# NASQAR: A web-based platform for high-throughput sequencing data analysis and visualization

**DOI:** 10.1101/709980

**Authors:** Ayman Yousif, Nizar Drou, Jillian Rowe, Mohammed Khalfan, Kristin C Gunsalus

## Abstract

**Background:** As high-throughput sequencing applications continue to evolve, the rapid growth in quantity and variety of sequence-based data calls for the development of new software libraries and tools for data analysis and visualization. Often, effective use of these tools requires computational skills beyond those of many researchers. To ease this computational barrier, we have created a dynamic web-based platform, NASQAR (Nucleic Acid SeQuence Analysis Resource).

**Results:** NASQAR offers a collection of custom and publicly available open-source web applications that make extensive use of a variety of R packages to provide interactive data analysis and visualization. The platform is publicly accessible at http://nasqar.abudhabi.nyu.edu/. Open-source code is on GitHub at https://github.com/nasqar/NASQAR, and the system is also available as a Docker image at https://hub.docker.com/r/aymanm/nasqarall. NASQAR is a collaboration between the core bioinformatics teams of the NYU Abu Dhabi and NYU New York Centers for Genomics and Systems Biology.

**Conclusions:** NASQAR empowers non-programming experts with a versatile and intuitive toolbox to easily and efficiently explore, analyze, and visualize their Transcriptomics data interactively. Popular tools for a variety of applications are currently available, including Transcriptome Data Preprocessing, RNA-seq Analysis (including Single-cell RNA-seq), Metagenomics, and Gene Enrichment.

## Background

Genomic data has experienced tremendous growth in recent years due to the rapid advancement of Next Generation Sequencing (NGS) technologies [1, 2]. Common applications include transcriptome profiling; de novo genome sequencing; metagenomics; and mapping of genomic variation, transcription factor binding sites, chromatin modifications, chromatin accessibility, and 3D chromatin conformation. Single-cell versions of these (e.g. [3]) and newer methods — such as spatial transcriptomics (e.g. [4]), CRISPR-based screens (e.g. [5]), and multi-modal profiling (simultaneous quantification of proteins and mRNAs, e.g. [6]) — are rapidly proliferating as new technical innovations come on the scene (e.g. [7, 8]). As the volume of data and diversity of applications continue to grow, so does the number of software libraries and tools for the analysis and visualization of these datasets. Many of the available tools for genomic data analysis require computational experience and lack a graphical user interface (GUI), making them inaccessible to many researchers whose work depends on them. Some of the common challenges include:

- Knowledge and experience in various programming/scripting languages (R, Python, shell, etc.)
- Data munging: pre-processing and reformatting for use with specific tools
- Limited computational resources (cpu, memory, and disk storage)
- Installation of software packages and dependencies. Many required tasks can be time consuming and tedious due to issues such as satisfying software or hardware requirements and resolving software dependencies. In one study [9], almost half (49%) of the published omics software tools that were randomly surveyed were found to be “difficult to install”. Moreover, the rapid churn of operating system updates and hardware configurations contributes to the gradual decline of a tool’s impact, usability, and lifetime.
- Software tools developed by researchers within academia are usually less “user-friendly”, due to either a lack of development resources or a lack of expertise in best practices for software engineering, such as cross-platform compatibility and user interface design [9]. For example, many available R GUI based tools, while featuring very useful and diverse functionality, lack simple error handling and/or informative feedback. This can render the application unmanageable if users cannot easily identify and remedy the causes of such errors.

NASQAR (Nucleic Acid SeQuence Analysis Resource) is a web-based platform that wraps popular high-level analysis and visualization tools in an intuitive and appealing interface. This platform addresses the above challenges by offering the following:

- Utilization of software and interface design best practices to craft user-friendly and intuitive tools that are based on commonly used analysis packages. This is important in order to lower the entry barrier to standard bioinformatics analysis and visualization workflows, thus providing greater independence for researchers with little or no programming experience. The platform may be used for QC, exploratory analysis, or production of publication-ready data files (such as normalized counts data) and figures (PCA plots, heatmaps, dendograms, UMAP/t-SNE etc.)
- A scalable virtualization architecture that is relatively simple to deploy on a personal computer, an organization’s private/public web servers, or on the cloud (AWS, Microsoft Azure, Google Cloud, etc.). Virtualization allows for the abstraction of software and operating system dependencies, thus alleviating difficulties in installation for end users. The scalable design is advantageous when deploying the platform online for multiple concurrent users, either for public use or internal use within a research facility. It uses open-source packages, which is particularly desirable for academic research institutions.
- Modular design of analysis categories. By decoupling data preprocessing, RNA-seq analysis, and gene enrichment applications from each other, userscan leverage these functions independently, thus allowing a greater versatility of analysis steps than fully integrated workflows.

The NASQAR platform provides a highly accessible, scalable, and user-friendly framework for versatile data analysis, comprising a consolidated toolbox of publicly available open-source applications (curated and vetted for good value and design) and custom applications developed in-house. While many useful web-based bioinformatics applications are now available, most focus exclusively on one type of analysis or application (e.g. bulk or single-cell RNA-seq, metagenomics, etc.) A few examples – some of which are included in NASQAR – include START[10], DEApp [11], TCC-GUI[12], Shiny-seq[13], GENAVi, is-CellR[14], and Shaman[15]. Fully integrated end-to-end analysis workflows such as GENAVi [16] employ a variety of R packages and/or other tools to streamline consecutive sequence analysis tasks (e.g. from preprocessing all the way to gene enrichment). While often desirable, this approach also restricts the end-user from performing just one of the implemented functions (such as gene enrichment), which is particularly useful for datasets generated independently using other tools or by external collaborators. NASQAR takes a different approach, aiming instead to empower non-programming experts with a ‘‘Swiss army knife” to perform a variety of sequence analysis tasks on their own. These may be accessed either as independent units or sequentially, with convenient interfaces to commonly used R data analysis packages and functions. This flexible framework offers a model resource for the community that can be extended to a broader range of applications through further development and collaboration.

## Implementation

The architecture of the NASQAR web platform is illustrated in Figure 1. NASQAR has been deployed on a cluster of virtual machines and is publicly accessible at http://nasqar.abudhabi.nyu.edu/. Docker [17] and Swarm provide containerization and cluster management, and the Traefik reverse proxy / load balancer (https://traefik.io/) manages requests and maintains sticky user sessions, which is essential in hosting Shiny applications for concurrent users. The scalable design makes it relatively easy to increase dedicated resources simply by adding more nodes to the Docker Swarm cluster, and thus to flexibly accommodate growth in computational demand as new applications are deployed and the user base expands. In addition, the platform has been deployed on AWS Cloud with Kubernetes (http://www.nasqar.com).

**Figure 1.**
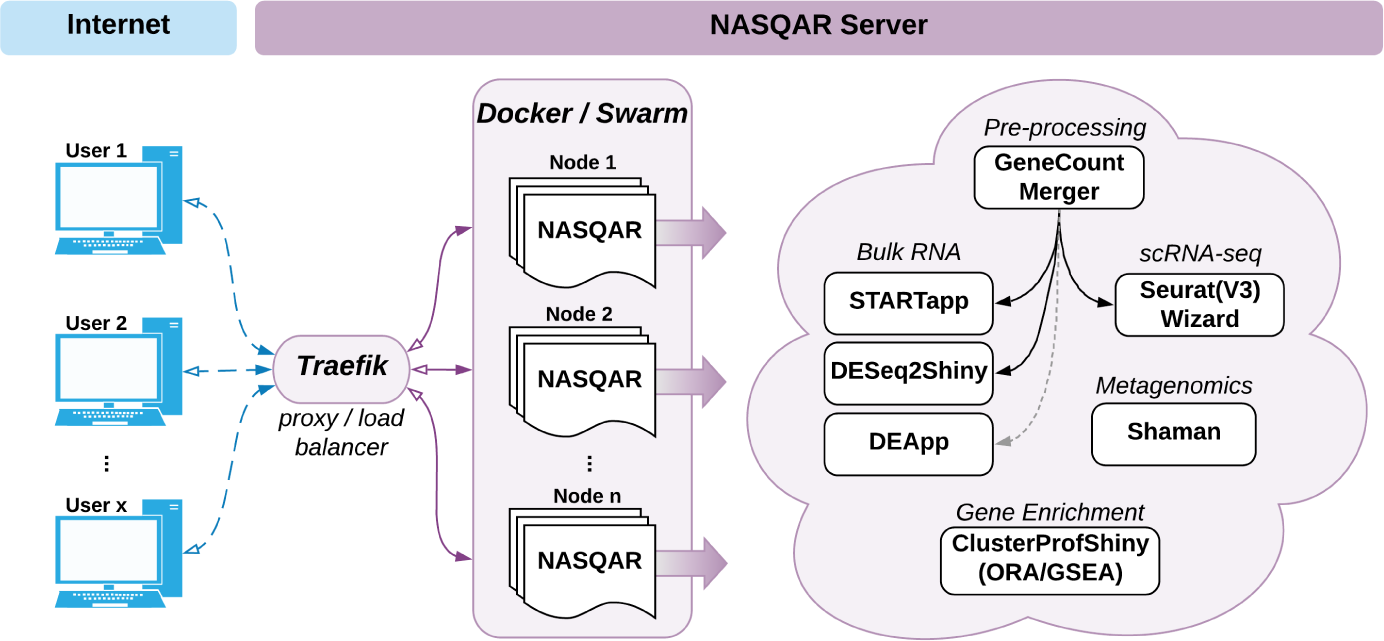
NASQAR Platform Architecture. A cluster of virtual machines at NYU Abu Dhabi serves NASQAR applications to multiple concurrent users. Applications are containerized and managed on the cluster using Docker and Swarm, while Traefik load-balances requests among available server nodes. Functionality includes merging gene counts, conversion of gene IDs to gene names, analysis of differential mRNA expression, metagenomics analysis, and functional enrichment analysis. Applications for bulk expression analysis include DESeq2, limma, and EdgeR. Single-cell RNAseq analysis with Seurat Wizards is built on top of the Seurat R package and includes options for filtering, normalization, dimensionality reduction (PCA), clustering, and t-SNE. Enrichment analysis includes applications for Gene Set Enrichment Analysis (GSEA) and Over-representation Analysis (ORA) built using the clusterProfiler R package.

A Docker image of NASQAR is publicly available through DockerHub and can be used to deploy the application seamlessly on any system, whether a local computer or a public or private internet server (such as a research institute’s intranet). Although data uploaded online for analysis with NASQAR (at http://nasqar.abudhabi.nyu.edu/) is by default discarded after a user’s session ends, this does not guarantee total data privacy. Where privacy is a concern (e.g. patient data), NASQAR may be deployed on either a restricted intranet or a personal computer. Moreover, using Docker allows deployment of the entire NASQAR toolbox with a one-time install, removing the hassle of having to manually satisfy the different software requirements of numerous individual applications. The source code is publicly available on GitHub and is actively maintained. Each individual application is hosted in its own GitHub repository and can be accessed and launched independently via R or R Studio. All applications have clear user guides with example data sets to help users get started and acclimate quickly. This is a major factor in improving usability and thus adoption of the tools.

NASQAR’s collection of applications is primarily implemented in R, a widely used and freely available statistical programming language [18]. Most of the analysis workflows are built using R libraries for genomics and computation. The frontend design employs several R libraries, such as Shiny[19], shinydashboard, shinyjs, shinyBS, shinycssloaders. These libraries and custom Javascript/CSS/HTML enhancements improve the user experience and overall usability with interface consistency, visual clarity, and ease of navigation. Familiar R packages used to build the applications include dplyr and tidyr for matrix data manipulation; ggplot2, heatmaply, and NMF for figure plotting; and BiocParallel for multi-threading support. Additional packages used in conjunction with specific tools are indicated below.

In addition to previously published software, we introduce several new applications we have developed that wrap around popular analysis packages, such as DESeq2 [20] and Seurat [21, 22] for bulk and single-cell RNA-seq analysis and visualization. Since most NASQAR applications require a matrix of gene counts as input, we have also built a convenient tool to assist with pre-processing, GeneCountMerger. Some of the applications provide a seamless transition from data pre-processing to downstream analysis. This implementation gives users the option of using multiple analysis applications without having to modify/reformat the input data set, thus allowing them to easily benchmark and compare the performance of different analysis software packages.

## Results and Discussion

NASQAR currently hosts tools for merging gene counts; conversion of gene IDs to gene names; and analysis of differential mRNA expression, gene function enrichment, and metagenomic profiling. Packages for bulk RNA-seq analysis include DESeq2, edgeR[23], and limma[24], while single-cell analysis is driven by Seurat (see Table 1 for a comprehensive overview of the applications). The Supplementary Materials include details on available applications along with example use cases. We believe the custom applications developed for NASQAR improve on several existing tools, as highlighted in the following application summaries.

**Table 1.**
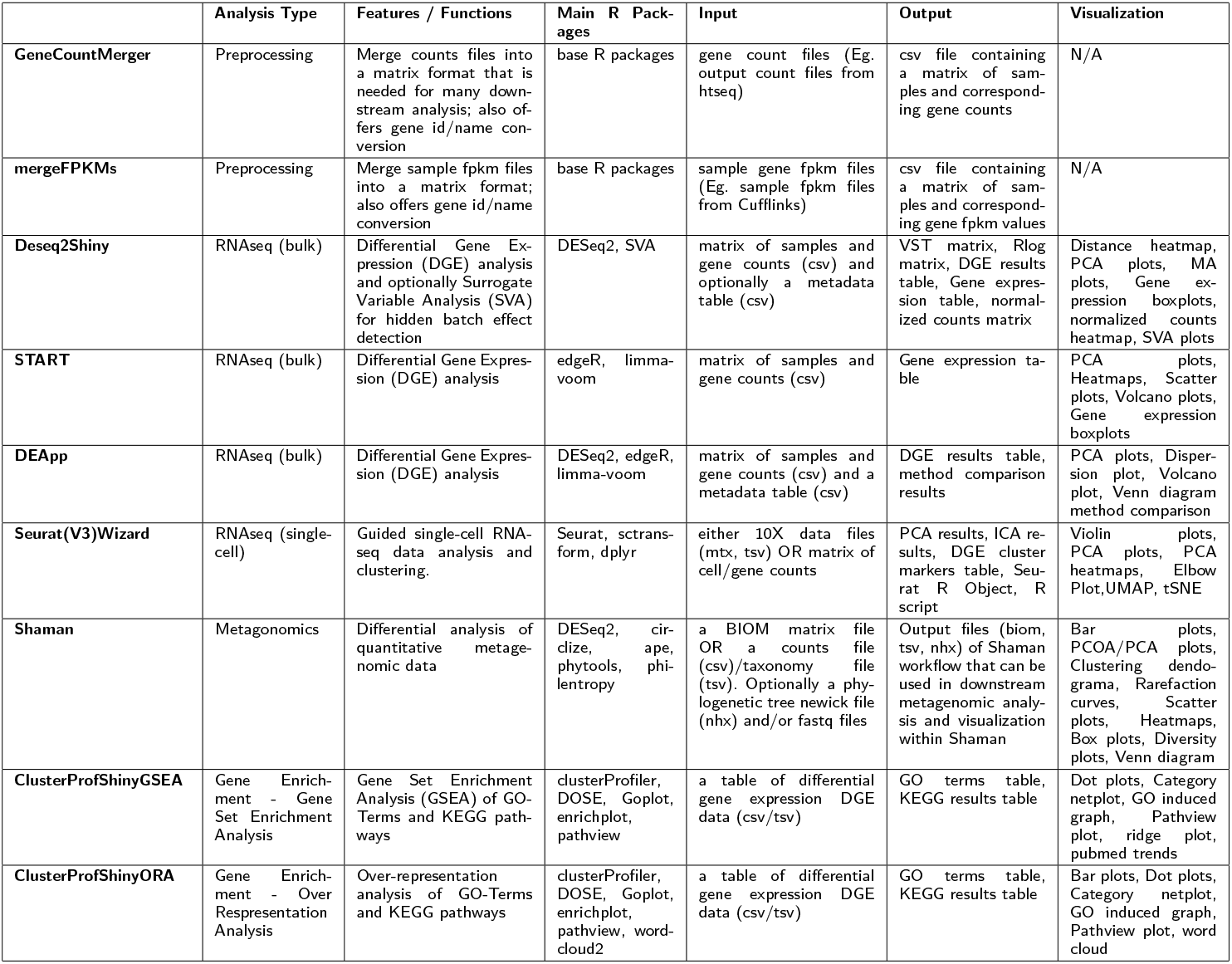
Comprehensive overview of NASQAR applications

### GeneCountMerger

This preprocessing tool is used to merge individual raw gene count files produced from software such as htseq-count [25] or featurecounts [26] (Figure 2). Options include:

- Merge individual sample count files into one matrix
- Merge multiple raw count matrices
- Convert Ensembl gene IDs to gene names
- Select from available genomes / versions
- Add pseudocounts
- Rename sample column headers
- Download merged counts file in .csv format
- Seamless transcriptome analysis following merging counts (Seurat Wizard for single-cell RNA analysis; DESeq2Shiny or START [10] for bulk RNA analysis)

**Figure 2.**
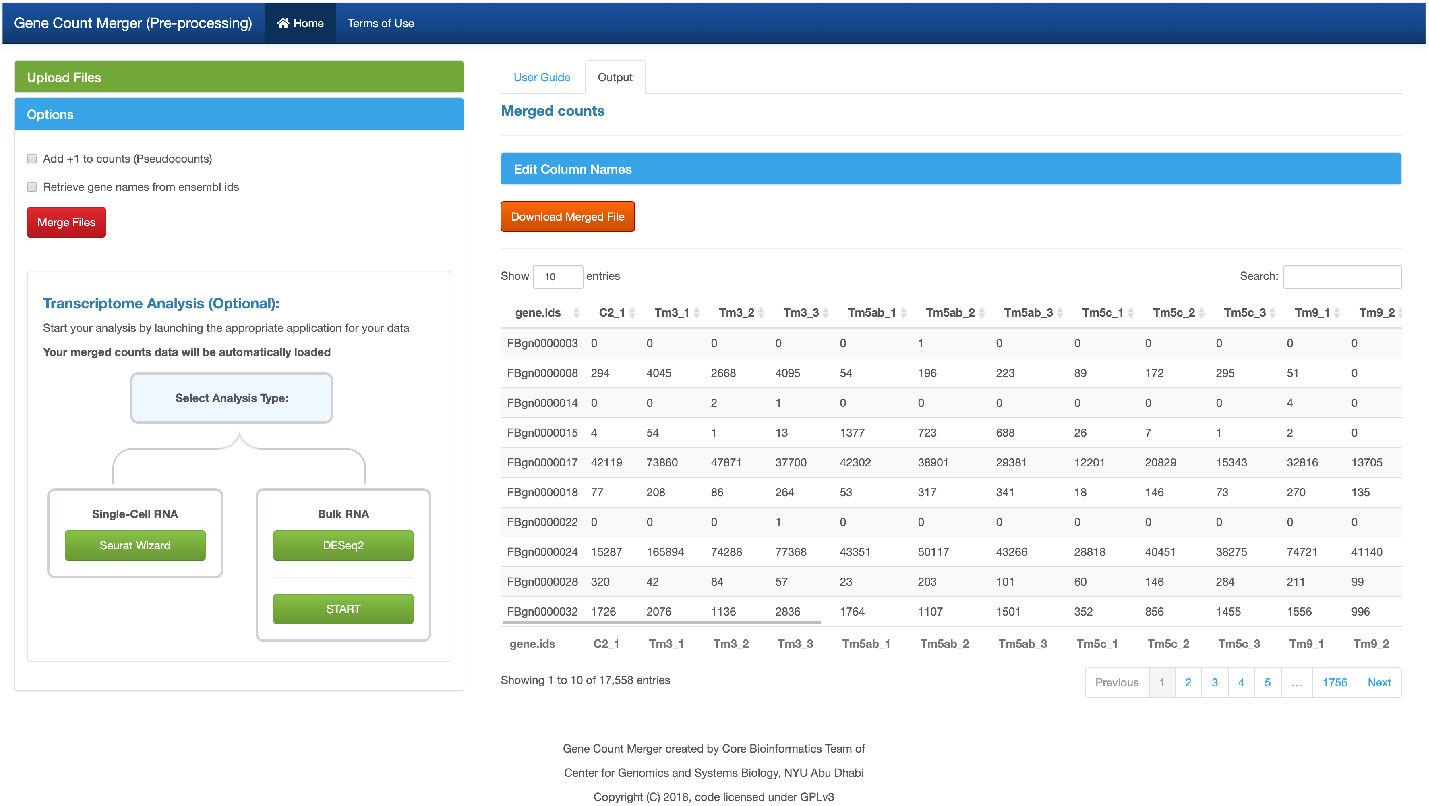
GeneCountMerger screenshot. A preprocessing utility to generate the gene count matrices required as input to many analysis tools. It can merge individual raw gene count files from htseq-count and other similar applications. Convenient features include conversion of Ensembl gene IDs to gene names for reference genomes and seamless launching of downstream analysis applications.

### Seurat Wizards

Seurat Wizards are wizard-style web-based interactive applications to perform guided single-cell RNA-seq data analysis and visualization using Seurat, a popular R package designed for QC, analysis, and exploration of single-cell RNAseq data (Figure 3). The design and implementation of the wizards offer an intuitive way to tune the analysis interactively by allowing users to inspect and visualize the output of intermediate steps and adjust parameters accordinly. In contrast, most web-based tools for scRNA-seq analysis, such as IS-CellR [14] and SCHNAPPs (https://c3bi-pasteur-fr.github.io/UTechSCB-SCHNAPPs/), provide integrated solutions that offer less opportunity for user intervention at intermediate steps. Some of the distinctive features of the wizards include, 1) allowing users to visually inspect the distribution of cells using violin plots and to select cutoff thresholds accordingly in order to filter out cells before starting the analysis, 2) Elbow/Jackstraw plots that assist the users in determining what dimensions to use for non-linear reduction. Both of these features can have significant consequences on downstream steps like clustering and differential analysis.

**Figure 3.**
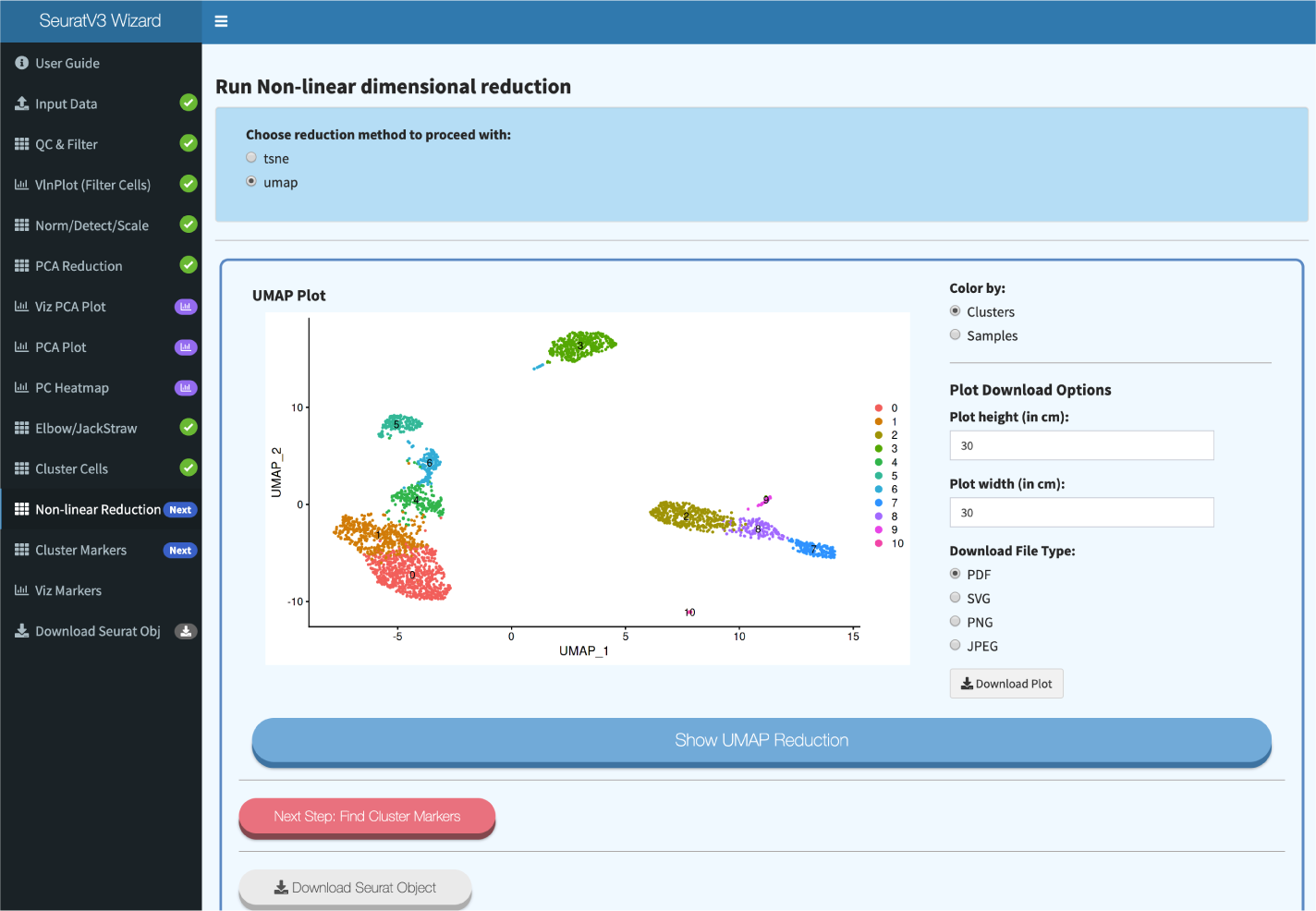
Seurat Wizard screenshot. Wizard-style web-based interactive applications based on Seurat, a popular R package designed for QC, analysis, and exploration of single-cell RNA-seq data. The wizards guide users through single-cell RNA-seq data analysis and visualization and provide an intuitive way to fine-tune parameters using feedback from results at each stage of the analysis. Functionality includes filtering, normalization, dimensionality reduction (PCA), clustering, and visualization with UMAP or t-SNE plots.

The Seurat Wizards follow closely the Seurat Guided Clustering Tutorials devised by the Seurat authors (https://satijalab.org/seurat/v3.0/pbmc3k_tutorial.html). Both Seurat versions 2 and 3 are currently supported. Users can follow the tutorials while using the Wizards and edit parameters at almost every step, which is instrumental in producing accurate results. Pre-processing (QC/filtering), normalization, dimensionality reduction, clustering (UMAP/t-SNE), and differential expression (cluster biomarkers) are all supported. To enhance the user experience and learning process, the wizards progress step-wise through the workflow. The workflow processing steps become available sequentially upon completion of each preceding task, thus avoiding visual clutter and focusing the user’s attention on the task at hand. One of the unique features of the Seurat Wizards is that they can accept as input either processed 10X Genomics data files or a matrix of gene counts, which eliminates the need for an additional pre-processing step. To address reproducibility, the last step of the wizard allows the user to download an R script with all of the R functions and parameters used for the analysis, along with the R object that contains all of the analyzed data, for further exploration in R/RStudio.

SeuratV3Wizard integrates several additional features like the UCSC Cell Browser (https://github.com/maximilianh/cellBrowser), which enables users to interactively visualize clusters and gene markers. Additional cell browser options will be implemented in future releases where feasible. It also includes the newly published sctransform method [27], which offers users the convenience of running an analysis using two slightly different workflows and comparing the results. We believe these differences in features and design give the Seurat Wizards more versatility and improve usability in comparison with other publicly available implementations.

### DESeq2Shiny

The DESeq2Shiny app is a Shiny wrapper around DESeq2, a popular R package for performing differential mRNA expression analysis of RNA-seq data (Figure 4). This web-based application provides functions for data normalization, transformation (e.g., rlog and vst for clustering), and estimation of dispersion and log fold-change. The results are all downloadable in csv format. Data visualizations include MA plots, heatmaps, dendograms, gene expression boxplots, and PCA.

**Figure 4.**
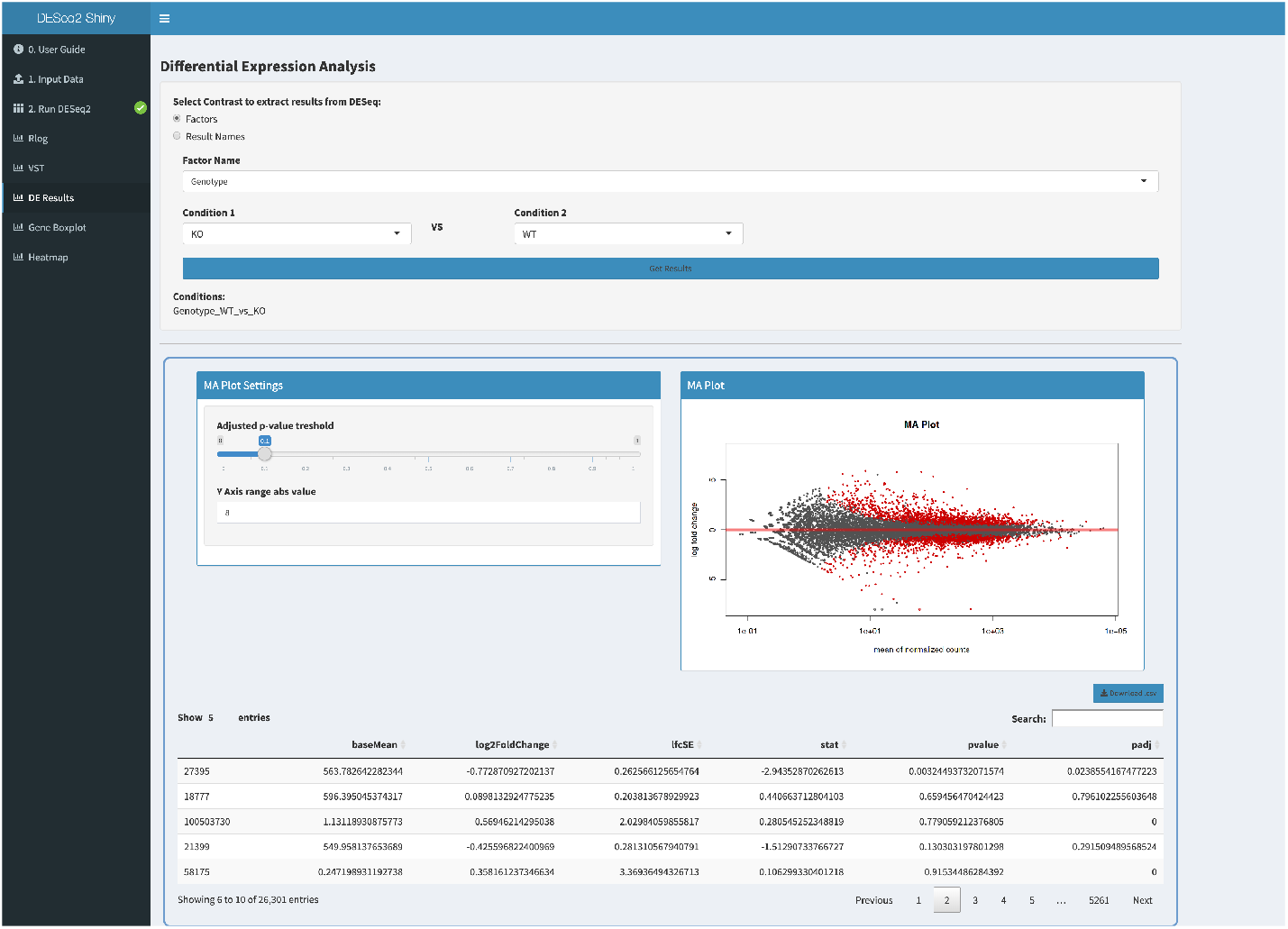
DESeq2Shiny screenshot. A web-based shiny wrapper around DESeq2, a popular R package for performing differential mRNA expression analysis of RNA-seq data.

The application is capable of working with simple experimental designs or complex experiments with multiple factors. For single-factor experiments with replicates, sample names can be parsed and grouped automatically given proper formatting. The experimental design table and formula can also be generated autonomously. For multifactor experiments, the table can be constructed easily within the “Edit Conditions” page, or an experiment design metadata (csv) file may be uploaded directly. The design formula expresses how the counts for each gene depend on the factor(s) and is editable within the “Edit Conditions” page. This gives users the option to specify experimental designs with multiple variables (e.g. *~group* + *condition)* and interaction terms (e.g. *~genotype* + *treatment* + *genotype*: *treatment).* In cases where no replicates exist, exploratory analysis (with no differential testing) may be performed by setting the formula to _-_1 (which signifies “no design”). Most other surveyed R Shiny applications for RNAseq differential gene expression (DGE) analysis (e.g. START, DEApp, TCC-GUI, and Shiny-seq) lack such flexible features. While users are warned that this type of analysis has low statistical significance and they should not draw strong conclusions based on the results, it is none the less implemented as a feature to allow for exploratory testing and hypothesis generation [28].

The DESeq2Shiny app interface design follows the same implementation as other apps on NASQAR: users are guided through the analysis, and subsequent steps become available when the current step is completed and valid. Users may also finetune analysis parameters interactively. This design, coupled with preloaded example datasets for single or multi-factor designs, aims to improve ease of use. Known batch effects can be modeled simply by adding the batch as a factor in the design matrix and formula. The application also offers hidden batch effect estimation using svaseq [29]. This allows for the estimation of surrogate variables, which can be included as adjustment factors in the design formula to correct subsequent downstream analysis.

### ClusterProfShiny

The ClusterProfilerShiny apps wrap the popular clusterProfiler [30] package, which implements methods to analyze and visualize functional profiles of genomic coordinates, genes, and gene clusters (Figure 5). Users can upload their own data from the output of DESeq2, for example, or import analyzed data from the upstream DESeq2Shiny app. These apps allow for quick and easy over-representation analysis (ORA) and gene set enrichment analysis (GSEA) of GO terms and KEGG pathways. Visuals produced include dot plots, word clouds, category net plots, enrichment map plots, GO induced graphs, GSEA plots, and enriched KEGG pathway plots using the Pathview [31] package.

**Figure 5.**
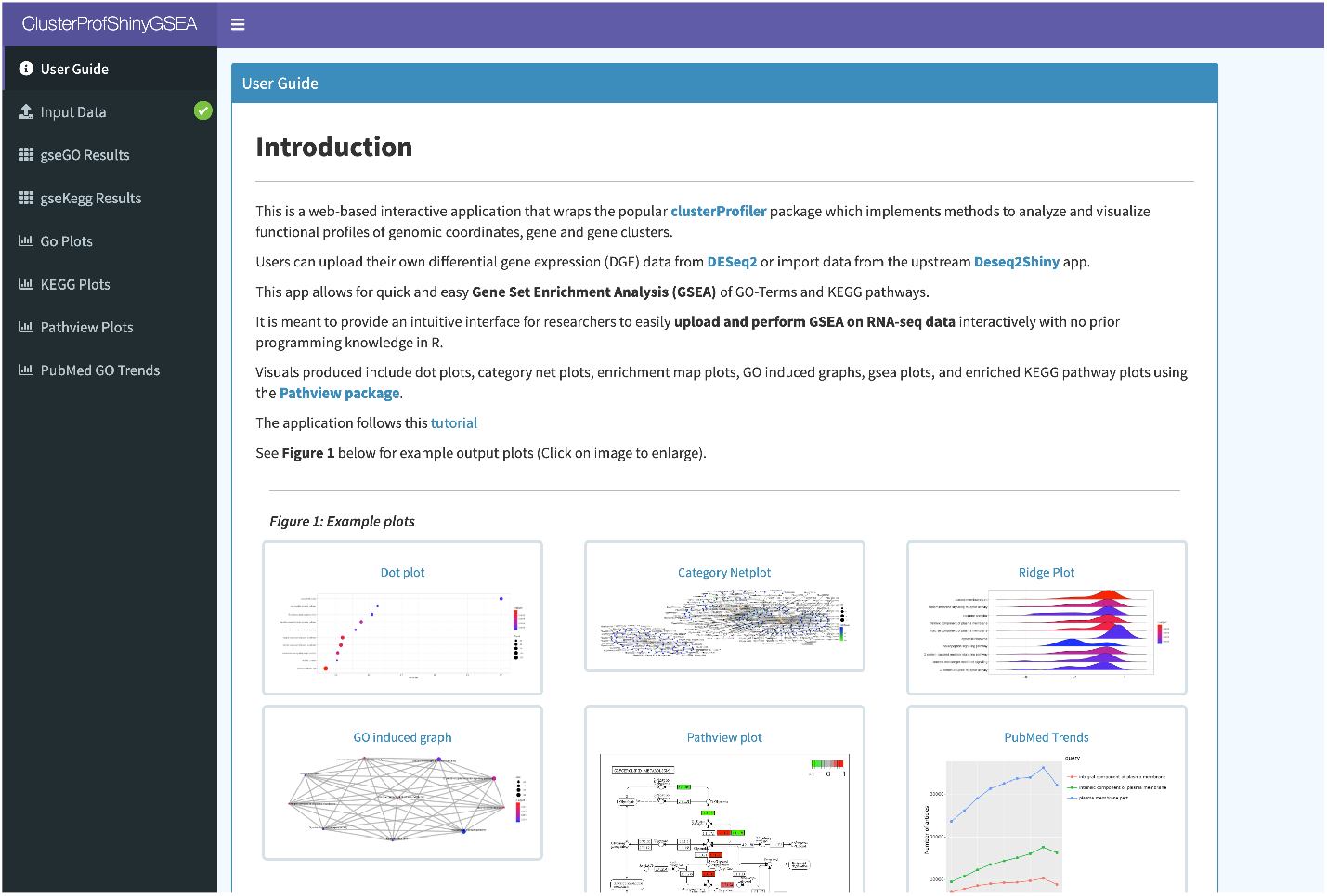
ClusterProfShinyGSEA screenshot. Web-based apps wrap the popular R package clusterProfiler for the analysis and visualization of functional themes and enrichment among gene clusters, using data from either DESeq2 or DESeq2Shiny. Both Gene Set Enrichment Analysis (GSEA) and Over-Representation Analysis (ORA) are implemented.

#### Other open-source apps

- **START**: a web-based RNA-seq analysis and visualization resource using edgeR and limma-voom. We have modified this application slightly from the published version to add options to some plots. We have also integrated it with GeneCountMerger so that once merging gene counts is complete, users may launch the START app and have their merged matrix data loaded automatically.
- **DEApp** [11]: an interactive web application for differential expression analysis using DESeq2, edgeR, limma-voom.
- **Shaman** [15]: a Shiny application that enables the identification of differentially abundant genera within metagenomic datasets. It wraps around the Generalized Linear Model implemented in DESeq2. It includes multiple visualizations and is compatible with common metagenomic file formats.

#### Ongoing and Future work

Numerous feature enhancements are planned or in progress to improve and expand functionality. For example, SeuratV3Wizard now provides the option to download an auto-generated R script and R object containing the executed code and results of a Seurat analysis. This enhances reproducibility by allowing users to inspect and document the specific commands and parameters used. Adding this option for other applications such as DESeq2shiny and ClusterProfiler(ORA/GSEA) will help users learn to understand their workflows in greater depth and will foster collaboration between experimental and computational biologists. In addition, we are continuously working to provide more online documentation for different use cases, to improve error handling for all NASQAR apps, and to evaluate possibilities for additional domain applications such as single-molecule long-read data. In order to facilitate broader deployment and ease of access, we have deployed NASQAR on AWS Cloud (available at www.nasqar.com). Cloud computing services open up opportunities for new analysis and visualization categories such as variant discovery, which requires both heavy computation and a large amount of data storage.

## Conclusion

The NASQAR platform offers a publicly available, comprehensive toolbox of interactive bioinformatics and visualization applications for sequence analysis that is accessible to all researchers with or without computational experience. NASQAR online services are currently deployed through NYU (with Docker/Swarm) and on AWS Cloud (with Kubernetes). In cases where data privacy is a major concern, the entire platform can be deployed privately on a personal computer or as a shared resource on a local intranet. Google Analytics traffic reports and GitHub activity show that the global user base is diverse and is increasing steadily, revealing rising demand in the community for easily accessible bioinformatics analysis and visualization platforms. NASQAR is under active development and will continue to offer user support and feature enhancements with future releases.

## Abbreviations

NASQAR: Nucleic Acid SeQuence Analysis Resource;
RNA-seq: RNA sequencing;
NGS: Next Generation Sequencing;
CRISPR: Clustered Regularly Interspaced ShortPalindromic Repeats;
GUI: Graphical User Interface;
PCA: Principal Components Analysis;
t-SNE: t-Distributed Stochastic Neighbor Embedding;
UMAP: Uniform Manifold Approximation and Projection;
QC: Quality Control;
UCSC: University of California, Santa Cruz;
rlog: regularized logarithm transformation;
vst: variance stabilizing transformation;
ORA: Over-Representation Analysis;
GSEA: Gene Set Enrichment Analysis;
GO: Gene Ontology;
KEGG: Kyoto Encyclopedia of Genes and Genomes;
START: Shiny Transcriptome Analysis Resource Tool;
Shaman: SHiny application for Metagenomic ANalysis;
csv: comma separated values

## Availability and requirements

**Project name:** NASQAR

**Project home page:**

**Operating system(s):** Platform independent

**Programming language:** R, JavaScript

**Other requirements:** Docker (version >= 17.03.0-ce)

**License:** GNU GPL.

**Any restrictions to use by non-academics:** none

## Declarations

### Ethics approval and consent to participate

Not applicable.

### Consent for publication

Not applicable.

### Availability of data and materials

NASQAR is publicly accessible at http://nasqar.abudhabi.nyu.edu/ and http://www.nasqar.com/. The platform is available as a Docker image at https://hub.docker.com/r/aymanm/nasqarall. NASQAR is open-source and the code is available through GitHub:

NASQAR (main page): https://github.com/nasqar/NASQAR

SeuratV3Wizard (scRNA): https://github.com/nasqar/seuratv3wizard

SeuratWizard (scRNA): https://github.com/nasqar/SeuratWizard

deseq2shiny (Bulk RNA): https://github.com/nasqar/deseq2shiny

GeneCountMerger (Pre-processing): https://github.com/nasqar/GeneCountMerger

ClusterProfShinyGSEA (Enrichment): https://github.com/nasqar/ClusterProfShinyGSEA

ClusterProfShinyORA (Enrichment): https://github.com/nasqar/ClusterProfShinyORA

### Competing interests

The authors declare that they have no competing interests.

### Funding

This work was supported by a grant from the NYU Abu Dhabi Research Institute to the NYU Abu Dhabi Center for Genomics and Systems Biology (CGSB).

### Author’s contributions

AY carried out the interface design and software development. ND defined platform requirements, contributed scripts and performed extensive software testing. JR contributed to the platform architecture design. MK contributed to the development of enrichment applications and provided guidance and extensive software testing. KCG supervised the project. All authors contributed to writing the manuscript. All authors approved the final version of the manuscript.

## Acknowledgements

The authors thank all the faculty and researchers in the NYU Abu Dhabi CGSB and Division of Biology for their excellent feedback, which has motivated the development of NASQAR. The authors would also like to acknowledge David Gresham and Siyu Sun (NYU New York CGSB) for their guidance during the development of the enrichment applications.

This research was carried out on the High Performance Computing resources at New York University Abu Dhabi. We extend special thanks to Fayizal Kunhi, NYU Abu Dhabi HPC.

## Author details

^1^NYU Abu Dhabi Center for Genomics & Systems Biology, Division of Biological Sciences, Abu Dhabi, United Arab Emirates. ^2^Center for Genomics & Systems Biology, Department of Biology, New York University, 10003 New York, United States.

## Additional Files

Additional file 1 — SupplementaryMaterials-NASQAR-final.pdf

This file includes supplementary materials such as instructions on how to launch

NASQAR and example use cases on data analysis and visualization.

## Notes

http://nasqar.abudhabi.nyu.edu/

### Competing Interest Statement

The authors have declared no competing interest.

### Summary of Updates

Minor edits to deseq2shiny section and Seurat Wizards section. Added a new table for a comprehensive summary of the applications included in NASQAR. Minor edit to supplementary materials

## References

1. Goodwin, S., McPherson, J.D., McCombie, W.R.: Coming of age: ten years of next-generation sequencing technologies. Nature Reviews Genetics 17, 333 (2016)

2. Wetterstrand, K.: DNA Sequencing Costs: Data from the NHGRI Genome Sequencing Program (GSP). Accessed on 07.08.2019. https://www.genome.gov/sequencingcostsdata

3. Zheng, M., Tian, S.Z., Capurso, D., Kim, M., Maurya, R., Lee, B., Piecuch, E., Gong, L., Zhu, J.J., Li, Z., Wong, C.H., Ngan, C.Y., Wang, P., Ruan, X., Wei, C.-L., Ruan, Y.: Multiplex chromatin interactions with single-molecule precision. Nature 566(7745), 558–562 (2019)

4. Stahl, P.L., Salmen, F., Vickovic, S., Lundmark, A., Navarro, J.F., Magnusson, J., Giacomello, S., Asp, M., Westholm, J.O., Huss, M., Mollbrink, A., Linnarsson, S., Codeluppi, S., Borg, Å., Ponten, F., Costea, P.I., Sahleen, P., Mulder, J., Bergmann, O., Lundeberg, J., Friseen, J.: Visualization and analysis of gene expression in tissue sections by spatial transcriptomics. Science 353(6294), 78–82 (2016)

5. Canver, M.C., Haeussler, M., Bauer, D.E., Orkin, S.H., Sanjana, N.E., Shalem, O., Yuan, G.-C., Zhang, F., Concordet, J.-P., Pinello, L.: Integrated design, execution, and analysis of arrayed and pooled crispr genome-editing experiments. Nature Protocols 13, 946 (2018)

6. Stoeckius, M., Hafemeister, C., Stephenson, W., Houck-Loomis, B., Chattopadhyay, P.K., Swerdlow, H., Satija, R., Smibert, P.: Simultaneous epitope and transcriptome measurement in single cells. Nature Methods 14, 865 (2017)

7. Stuart, T., Satija, R.: Integrative single-cell analysis. Nature Reviews Genetics 20(5), 257–272 (2019)

8. Mimitou, E.P., Cheng, A., Montalbano, A., Hao, S., Stoeckius, M., Legut, M., Roush, T., Herrera, A., Papalexi, E., Ouyang, Z., Satija, R., Sanjana, N.E., Koralov, S.B., Smibert, P.: Multiplexed detection of proteins, transcriptomes, clonotypes and crispr perturbations in single cells. Nature Methods 16(5), 409–412 (2019)

9. Mangul, S., Martin, L.S., Eskin, E., Blekhman, R.: Improving the usability and archival stability of bioinformatics software. Genome Biology 20(1), 47 (2019)

10. Sklenar, J., Nelson, J.W., Minnier, J., Barnes, A.P.: The START App: a web-based RNAseq analysis and visualization resource. Bioinformatics 33(3), 447–449 (2016)

11. Li, Y., Andrade, J.: Deapp: an interactive web interface for differential expression analysis of next generation sequence data. Source Code for Biology and Medicine 12(1), 2 (2017)

12. Su, W., Sun, J., Shimizu, K., Kadota, K.: Tcc-gui: a shiny-based application for differential expression analysis of rna-seq count data. BMC Research Notes 12(1), 133 (2019). doi:10.1186/s13104-019-4179-2

13. Sundararajan, Z., Knoll, R., Hombach, P., Becker, M., Schultze, J.L., Ulas, T.: Shiny-seq: advanced guided transcriptome analysis. BMC Research Notes 12(1), 432 (2019). doi:10.1186/s13104-019-4471-1

14. Patel, M.V.: iS-CellR: a user-friendly tool for analyzing and visualizing single-cell RNA sequencing data. Bioinformatics 34(24), 4305–4306 (2018)

15. Quereda, J.J., Dussurget, O., Nahori, M.-A., Ghozlane, A., Volant, S., Dillies, M.-A., Regnault, B., Kennedy, S., Mondot, S., Villoing, B., Cossart, P., Pizarro-Cerda, J.: Bacteriocin from epidemic listeria strains alters the host intestinal microbiota to favor infection. Proceedings of the National Academy of Sciences 113(20), 5706–5711 (2016)

16. Reyes, A.L.P., Silva, T.C., Coetzee, S.G., Plummer, J.T., Davis, B.D., Chen, S., Hazelett, D.J., Lawrenson, K., Berman, B.P., Gayther, S.A., Jones, M.R.: Genavi: a shiny web application for gene expression normalization, analysis and visualization. BMC Genomics 20(1), 745 (2019). doi:10.1186/s12864-019-6073-7

17. Merkel, D.: Docker: Lightweight linux containers for consistent development and deployment. Linux J. 2014(239) (2014)

18. R Core Team: R: A Language and Environment for Statistical Computing. R Foundation for Statistical Computing, Vienna, Austria (2017). R Foundation for Statistical Computing. https://www.R-project.org/

19. Chang, W., Cheng, J., Allaire, J., Xie, Y., McPherson, J.: Shiny: Web Application Framework for R. (2018). R package version 1.1.0. https://CRAN.R-project.org/package=shiny

20. Love, M.I., Huber, W., Anders, S.: Moderated estimation of fold change and dispersion for rna-seq data with deseq2. Genome Biology 15, 550 (2014)

21. Butler, A., Hoffman, P., Smibert, P., Papalexi, E., Satija, R.: Integrating single-cell transcriptomic data across different conditions, technologies, and species. Nature Biotechnology 36, 411 (2018)

22. Stuart, T., Butler, A., Hoffman, P., Hafemeister, C., Papalexi, E., Mauck, W.M.I., Hao, Y., Stoeckius, M., Smibert, P., Satija, R.: Comprehensive integration of single-cell data. Cell 177(7), 1888–1902 (2019)

23. Robinson, M.D., McCarthy, D.J., Smyth, G.K.: edgeR: a Bioconductor package for differential expression analysis of digital gene expression data. Bioinformatics 26(1), 139–140 (2009). doi:10.1093/bioinformatics/btp616. http://oup.prod.sis.lan/bioinformatics/article-pdf/26/1/139/443156/btp616.pdf

24. Ritchie, M.E., Phipson, B., Wu, D., Hu, Y., Law, C.W., Shi, W., Smyth, G.K.: limma powers differential expression analyses for RNA-sequencing and microarray studies. Nucleic Acids Research 43(7), 47–47 (2015). doi:10.1093/nar/gkv007. http://oup.prod.sis.lan/nar/article-pdf/43/7/e47/7207289/gkv007.pdf

25. Anders, S., Pyl, P.T., Huber, W.: HTSeq—a Python framework to work with high-throughput sequencing data. Bioinformatics 31(2), 166–169 (2014)

26. Smyth, G.K., Shi, W., Liao, Y.: featureCounts: an efficient general purpose program for assigning sequence reads to genomic features. Bioinformatics 30(7), 923–930 (2013)

27. Hafemeister, C., Satija, R.: Normalization and variance stabilization of single-cell rna-seq data using regularized negative binomial regression. bioRxiv (2019)

28. Anders, S., Huber, W.: Differential expression analysis for sequence count data. Genome Biology 11(10), 106 (2010). doi:10.1186/gb-2010-11-10-r106

29. Leek, J.T.: svaseq: removing batch effects and other unwanted noise from sequencing data. Nucleic Acids Research 42(21), 161–161 (2014). doi:10.1093/nar/gku864. http://oup.prod.sis.lan/nar/article-pdf/42/21/e161/9479130/gku864.pdf

30. Yu, G., Wang, L.-G., Han, Y., He, Q.-Y.: clusterprofiler: an r package for comparing biological themes among gene clusters. OMICS: A Journal of Integrative Biology 16(5), 284–287 (2012)

31. Luo, W., Brouwer, C.: Pathview: an R/Bioconductor package for pathway-based data integration and visualization. Bioinformatics 29(14), 1830–1831 (2013)

