## Supplementary Materials for "NASQAR: A web-based platform for high-throughput sequencing data analysis and visualization"

### 1) NASQAR platform and implementation details:

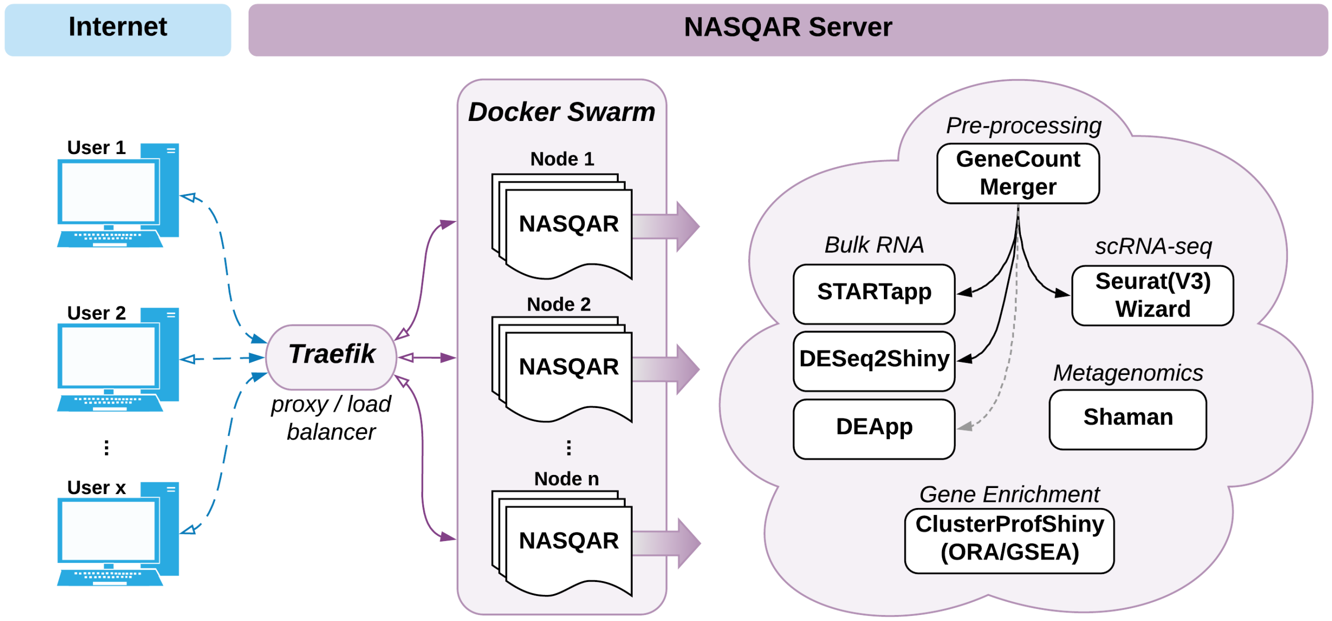


**Fig. 1.** NASQAR Platform Architecture. A cluster of virtual machines at NYU Abu Dhabi serves NASQAR applications to multiple concurrent users. Applications are containerized and managed on the cluster using Docker and Swarm, while Traefik load-balances requests among available server nodes. Functionality includes merging gene counts, conversion of gene IDs to gene names, analysis of differential mRNA expression, metagenomics analysis, and gene set and functional enrichment analysis. Applications for bulk expression analysis include DESeq2, limma, and EdgeR. Single-cell RNAseq analysis with Seurat Wizards is built on top of the Seurat R package and includes options for filtering, normalization, dimensionality reduction (PCA), clustering, and t-SNE. Enrichment analysis includes applications for Gene Set Enrichment Analysis (GSEA) and Over-representation Analysis (ORA) built using the clusterProfiler R package.

The architectural framework of the NASQAR web platform is illustrated in Figure 1. NASQAR has been deployed on a cluster of virtual machines and is publicly accessible at http://nasqar.abudhabi.nyu.edu/. Docker (Merkel 2014) and Swarm (Soppelsa and Kaewkasi 2017) provide containerization and cluster management, and the Traefik reverse proxy / load balancer (https://traefik.io/) manages requests and maintains sticky user sessions, which is essential for hosting Shiny applications. This framework allows access to multiple users concurrently while providing sufficient resources (RAM/CPU) for the applications. In anticipation of growing computational demand and the addition of more applications, the scalable design makes it relatively easy to increase dedicated resources simply by adding more nodes to the cluster.

A Docker image of NASQAR is publicly available through DockerHub and can be used to deploy the application seamlessly on any system with Docker installed, whether a local computer or a public server. In addition, each application can be installed and launched on its own, saving users from the hassle of satisfying the different software and hardware requirements. The source code is available publicly on GitHub and is actively maintained. All applications have clear user guides with example data sets to help users get started and acclimate quickly.

NASQAR comprises a collection of applications primarily implemented in R, a widely used and freely available statistical programming language (R Core Team 2017). Most of the analysis workflows are built using R libraries for genomics and computation. The front-end design utilizes the R Shiny (Chang *et al.* 2018) library and is supported by JavaScript/CSS to enhance usability and improve overall user experience.

In addition to previously published software, we introduce here several new applications we have developed that wrap around popular analysis packages, such as DESeq2 (Love *et al.* 2014) and Seurat (Butler *et al.* 2018) for bulk and single-cell RNA-seq analysis and visualization, respectively. Since most of the analysis applications in NASQAR require a matrix of gene counts as input, we have also built a convenient tool to assist with preprocessing, GeneCountMerger. Some of the applications have been integrated to provide a seamless transition from data preprocessing to downstream analysis. This implementation gives users the option of using multiple analysis applications without having to modify/reformat the input data set, thus allowing them to easily benchmark and compare the performance of different analysis software packages.

The following is a description of each application currently hosted by NASQAR:

#### *1.1 GeneCountMerger*

This preprocessing tool is used to merge individual raw gene count files produced from software such as htseq-count (Pyl *et al.* 2014) or featurecounts (Smyth *et al.* 2013). Options include:

- Merge individual sample count files into one matrix _
- Merge multiple raw count matrices _
- Convert Ensembl gene IDs to gene names _
- Select from available genomes / versions _
- Add pseudocounts _
- Rename sample column headers _
- Download merged counts file in .csv format _
- Seamless transcriptome analysis following count merger (Seurat _Wizard for single-cell RNA analysis; DESeq2Shiny or START (Sklenar *et al.* 2016) for bulk RNA analysis) _

#### *1.2 Seurat Wizards*

Seurat Wizards are wizard-style web-based interactive applications to perform guided single-cell RNA-seq data analysis and visualization. They are based on Seurat, a popular R package designed for QC, analysis, and exploration of single-cell RNAseq data. The wizard style makes it intuitive to go back and forth between steps and adjust parameters based on the results/feedback of different

outputs/plots/steps, giving the user the ability to interactively tune the analysis. SeuratWizard and SeuratV3Wizard implementations provide support for Seurat versions 2 and 3 (Stuart *et al.* 2019), respectively. _
Another web-based tool for scRNA-seq analysis, IS-CellR(Patel 2018), has recently been described that also utilizes Seurat v2. The SeuratWizard and SeuratV3Wizard take a different approach to design and implementation and follow closely the Seurat Guided Clustering Tutorials devised by the authors (https://satijalab.org/seurat/v3. 0/pbmc3k_tutorial.html). Users can follow the tutorials while using the Wizards and can edit parameters at almost every step, which is instrumental in producing accurate results. A unique feature of the Seurat Wizards is that they can accept as input processed 10X Genomics data files in place of a matrix of gene counts, which eliminates the need for this additional pre-processing step. SeuratV3Wizard integrates several additional features like the UCSC Cell Browser (https://github. com/maximilianh/cellBrowser), enabling users to interactively visualize clusters and gene markers, and the newly published sctransform method (Hafemeister and Satija 2019), which gives users the ability to run the analysis using two slightly different workflows and compare the results. These differences in features and design give the Seurat Wizards more versatility and improve usability in comparison with other publicly available implementations of Seurat.

#### *1.3 Deseq2Shiny*

The Deseq2Shiny app is a Shiny wrapper around DESeq2, a popular R package for performing differential mRNA expression analysis of RNA-seq data. This web-based application implements the standard default workflow outlined in_the DESeq2 Bioconductor tutorial (https://bioconductor.org/ packages/devel/bioc/vignettes/DESeq2/inst/doc/DESeq2. html). This includes normalization, data transformation (e.g., rlog and _vst for clustering), and estimation for dispersion and log fold change. This app follows the same implementation as other apps on NASQAR, whereby users can fine tune the analysis parameters interactively.

#### *1.4 ClusterProfilerShiny*

The ClusterProfilerShiny apps wrap the popular clusterProfiler (Yu *et al.* 2012) package, which implements methods to analyze and visualize functional profiles of genomic coordinates, genes, and gene clusters. Users can upload their own data from DESeq2 or import data from the upstream Deseq2Shiny app. These apps allow for quick and easy over-representation analysis (ORA) and gene set enrichment analysis (GSEA) of GO terms and KEGG pathways. Visuals produced include dot plots, word clouds, category net plots, enrichment map plots, GO induced graphs, GSEA plots, and enriched KEGG pathway plots using the Pathview package (Luo and Brouwer 2013).

#### *1.5 Other open-source apps*

- START: a web-based RNA-seq analysis and visualization resource. We have modified this application slightly from the open-source version to add options to some plots. We have also integrated it with GeneCountMerger so that once merging gene counts is complete, users can launch the START app and have their merged matrix data loaded automatically. _
- DEApp (Li and Andrade 2017): an interactive web application for differential expression analysis. _

- Shaman (Quereda *et al.* 2016): a Shiny application that enables the identification of differentially abundant genera within metagenomic datasets. It wraps around the Generalized Linear Model implemented in DESeq2. It includes multiple visualizations, and is compatible with common metagenomic file formats. _

### 2) Launch NASQAR using Docker (Recommended):

The recommended way to get NASQAR running is using Docker. The reason is that applications hosted within NASQAR have many package dependencies (R and OS) that might be tedious and very time consuming for the average user especially when trying to support different OS's (Windows/Linux/OSX). **Prerequisite**: Make sure Docker (version >= 17.03.0-ce, https://docs.docker.com/install/ ) is installed.

Run NASQAR docker image as follows:

a) docker run -p 80:80 aymanm/nasqarall:nasqar (runs on port 80)
   If you are running this in your personal laptop/PC, access NASQAR using a modern web browser at the following URL http://localhost/
b) docker run -p 8083:80 aymanm/nasqarall:nasqar (runs on port 8083)
   If you are running this in your personal laptop/PC, access NASQAR using a modern web browser at the following URL http://localhost:8083/
c) If you are hosting this as a service at your organization, make sure the specified port is not blocked with a firewall so users can access the service. Execute the same commands as a) and b). Users can access NASQAR using a modern web browser at the following URL http://<server_ip>:port/

**Note**: All apps in NASQAR can be launched individually. Visit each app's GitHub page for relevant instructions. For 3rd party apps hosted on NASQAR, please refer to their Github repositories.

**App GitHub pages:**

- SeuratV3Wizard (scRNA): https://github.com/nasqar/seuratv3wizard
- SeuratWizard (scRNA): https://github.com/nasqar/SeuratWizard
- deseq2shiny (Bulk RNA): https://github.com/nasqar/deseq2shiny
- GeneCountMerger (Pre-processing): https://github.com/nasqar/GeneCountMerger
- ClusterProfShinyGSEA (Enrichment): https://github.com/nasqar/ClusterProfShinyGSEA
- ClusterProfShinyORA (Enrichment): https://github.com/nasqar/ClusterProfShinyORA
- NASQAR (main page): https://github.com/nasqar/NASQAR

### 3) Example Use Case 1 (DGE and GSEA):

In this example we will show, using only your web browser, how you can start with individual sample gene count files (eg output of htseq counts) and carry out Differential Gene Expression (DGE) and Gene Set Enrichment analysis using DESeq2 and clusterProfiler R packages respectively.
The datasets provided in this example use case have been download from the ENSEMBL expression ATLAS (https://www.ebi.ac.uk/gxa/experiments/E-MTAB-970/Results). They belong to the ENSEMBL expression ATLAS experiment title "***Transcription profiling by high throughput sequencing of Sox17.Epi and Endo cells from mouse embryos***" ([dataset] 2016). The data comprises of 6 mouse RNA-seq samples, across two conditions. They were selected as an example dataset only, no other criteria were used in their selection.

#### *Step 1: Merge counts (GeneCountMerger):*

a) Download example count files zip from here (https://drive.google.com/file/d/1OB2vojHqscLAHZWGETHdGK0yaZ9r06PT/iew?usp=sharing )
b) Extract zip file
c) Launch GeneCountMerger http://nasqar.abudhabi.nyu.edu/GeneCountMerger/
d) Click browse and select all gene count files:

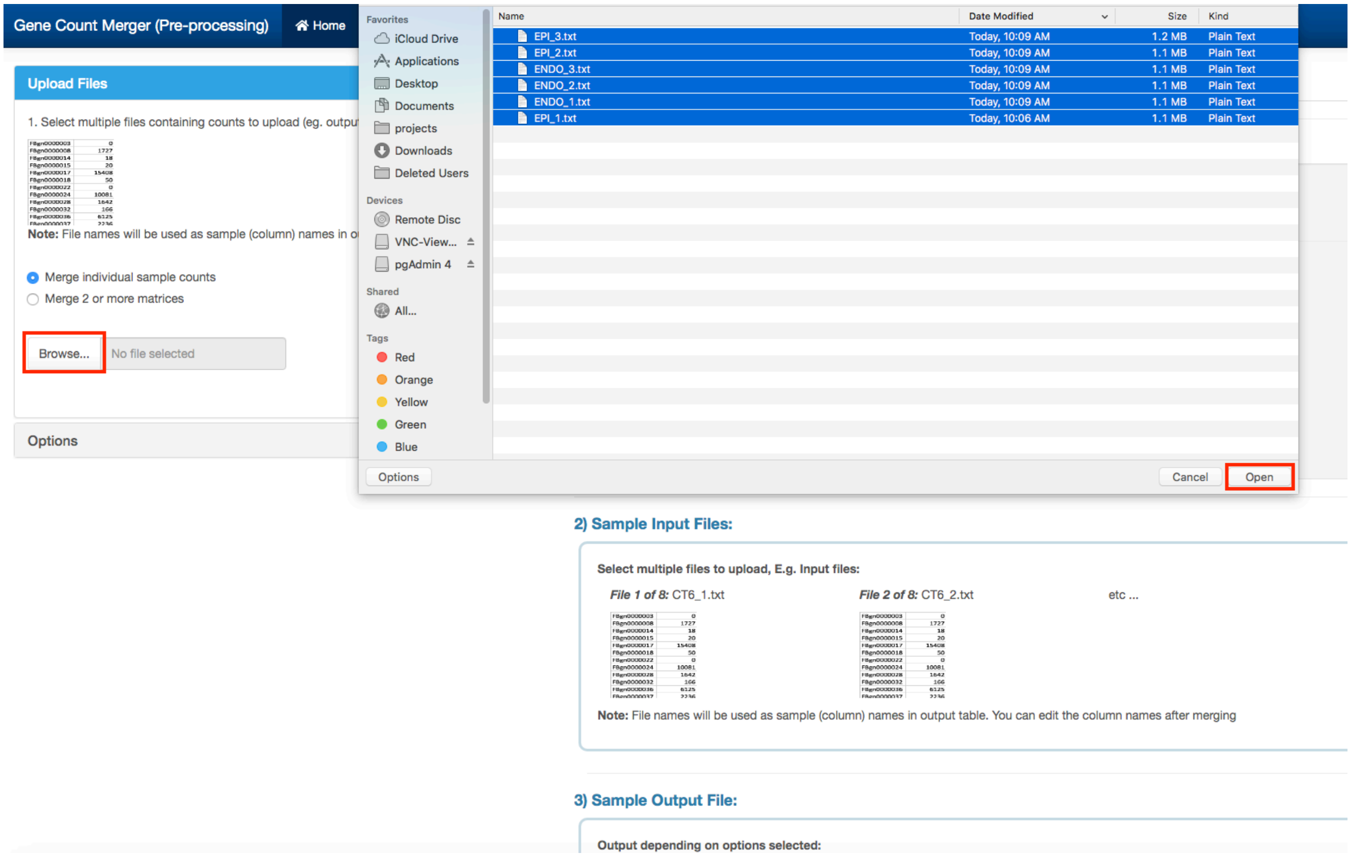

e) Once loaded, click the red Merge button. Now all counts files have been consolidated into one table. You can now download and save it as a .csv file

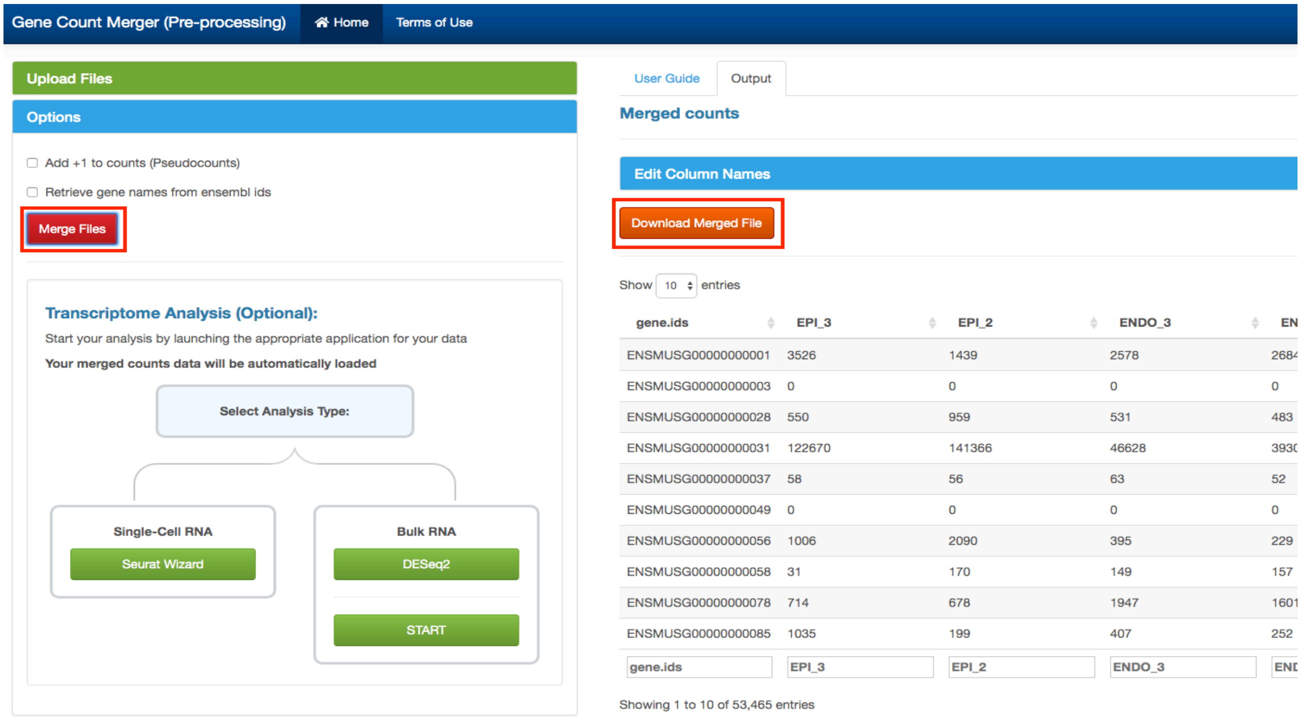


***Step 2: Differential Gene Expression analysis (Deseq2Shiny):***

a) Carrying on from the previous step, under “**Select Analysis Type**” select “**DESeq2**”

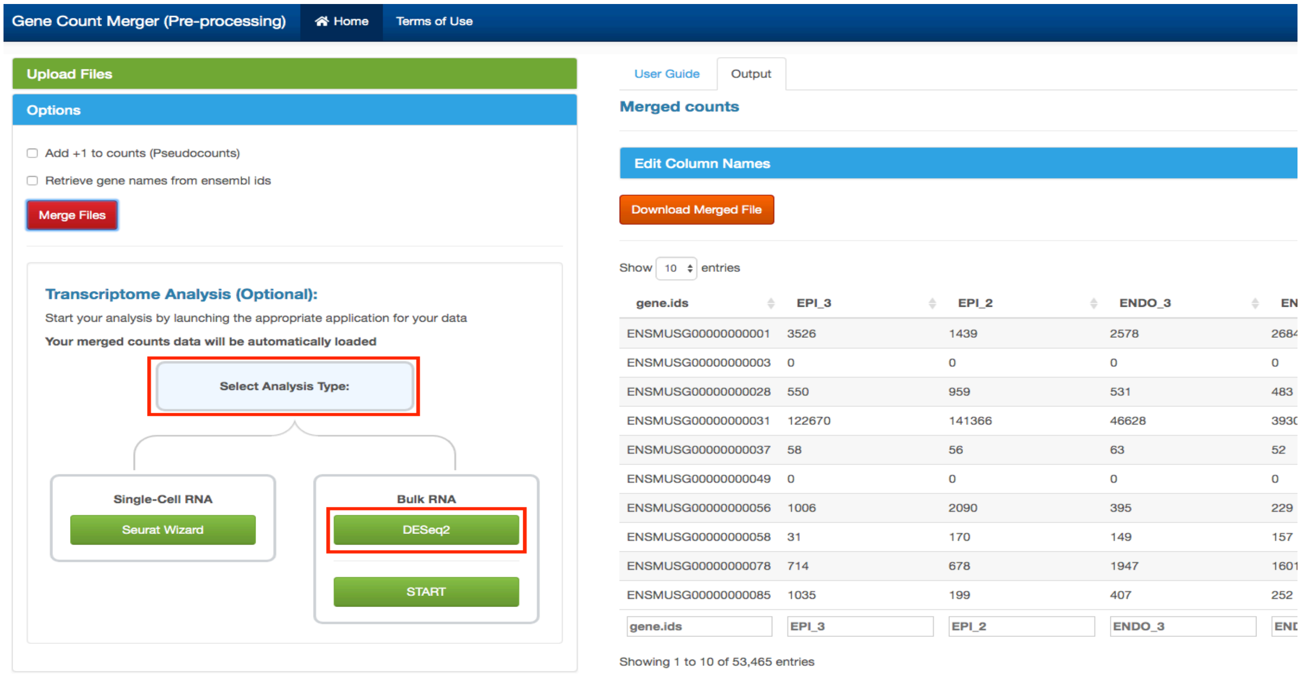


b) The Deseq2Shiny app will be launched in a new tab, with all counts data already loaded onto it. You can set the minimum number of counts to 1000 for example (this is mainly for speed up, you can continue with out it) and click “**Filter**”. Then click “**Next: Conditions**”

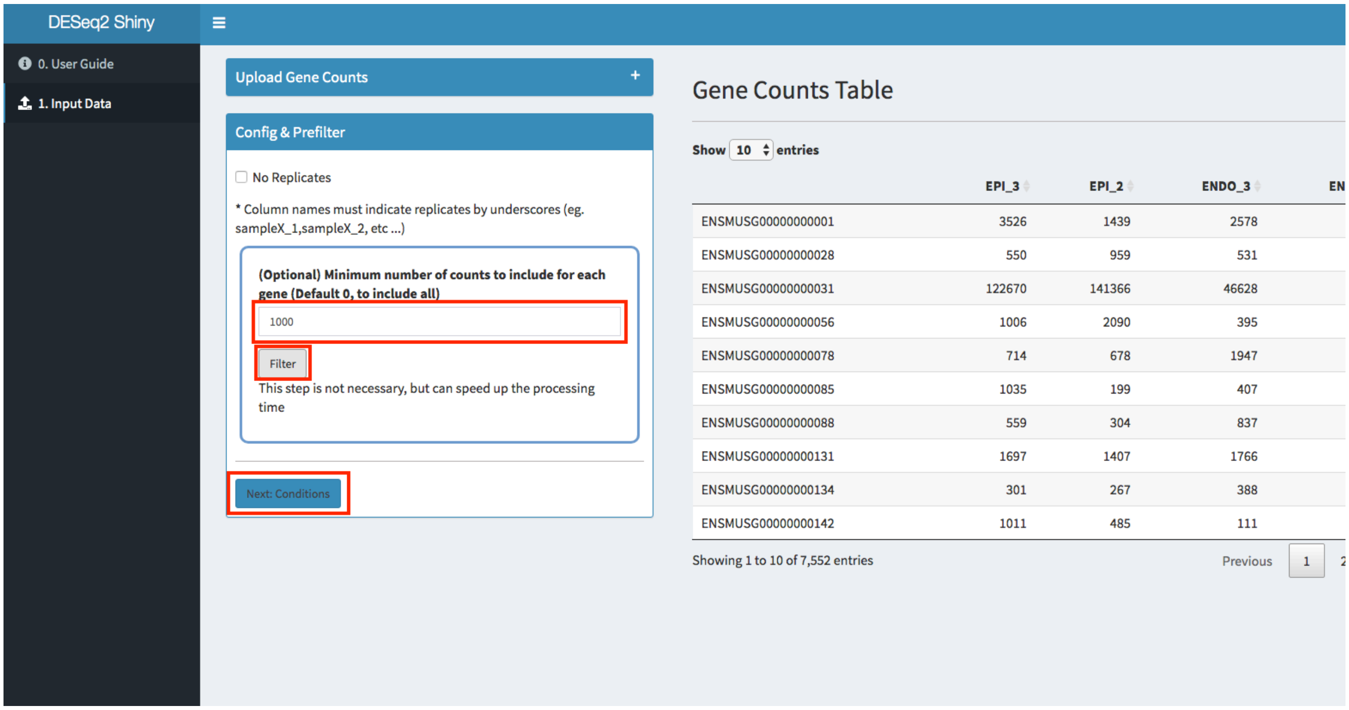


c) You can verify the sample/conditions table on the left, then click “**Run DESeq2**”. The replicate/condition IDs are inferred from the naming of the raw counts files. For example, ENDO_1 will be automatically determined to mean that the sample belongs to the condition

“ENDO”, and that it is the first replicate. In case a user supplies filenames that are named differently, this page will allow users to tag their replicates to the appropriate conditions.

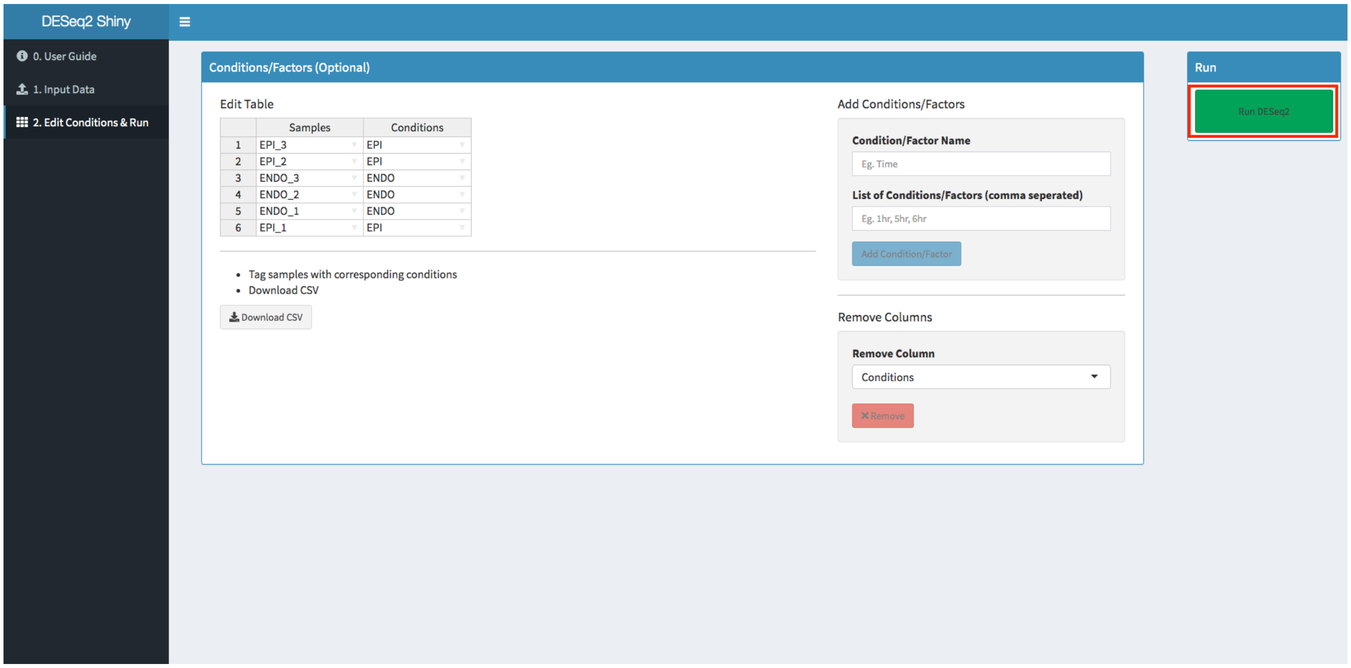


d) Once DESeq2 has completed, you will be able to see rlog/vst transformation matrices and PCA and distance heatmap plots.

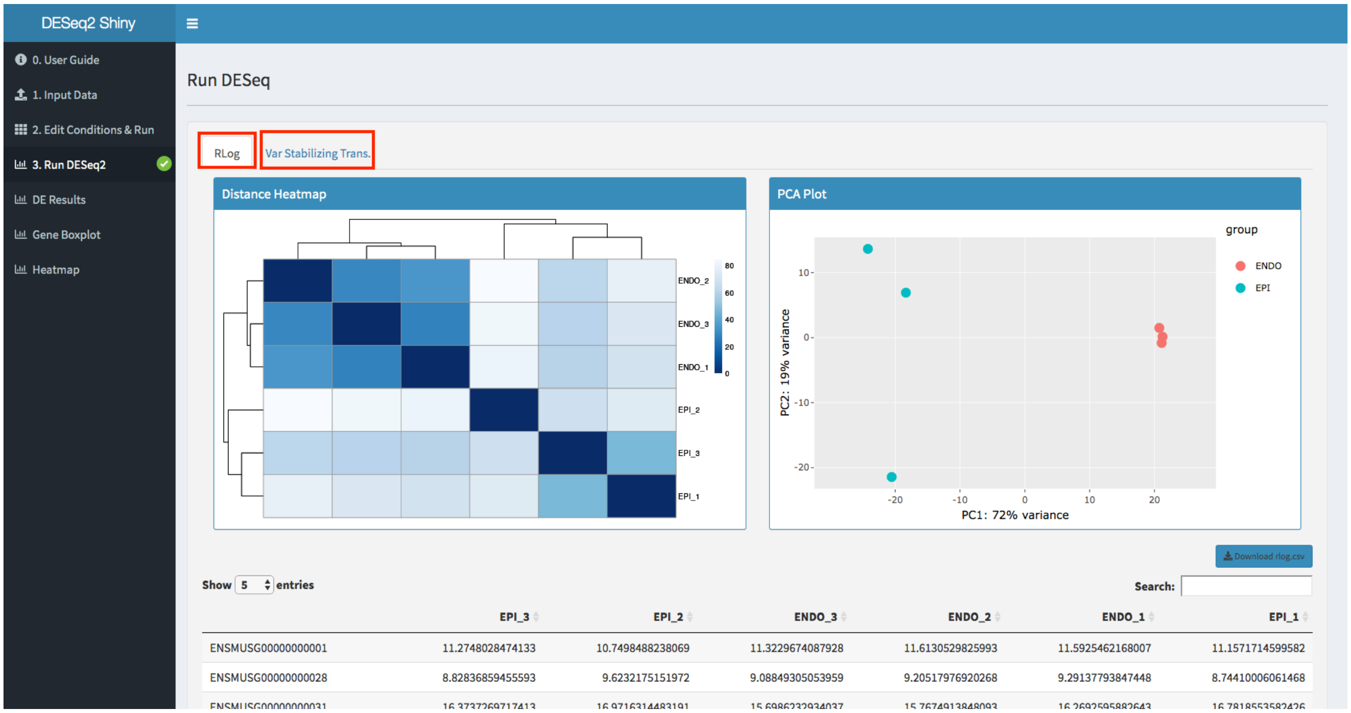

e) Next, go to “**DE Results**” to be able to run comparisons between different sample conditions. Select two different conditions, for example “EPI” and “ENDO” and then click “**Get Results**”

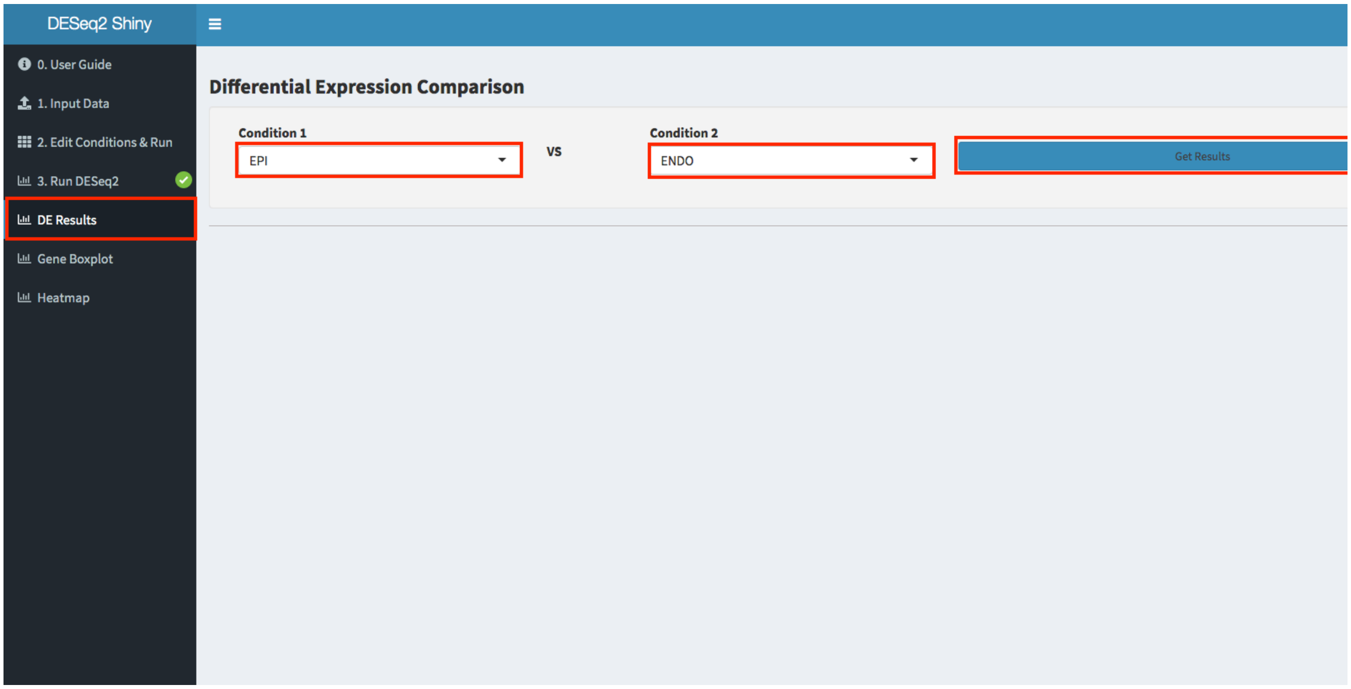


f) Scroll down and you can see the DE comparison results. Click “**Download .csv**”. This will save a CSV file locally called “EPI_vs_ENDO.csv”.

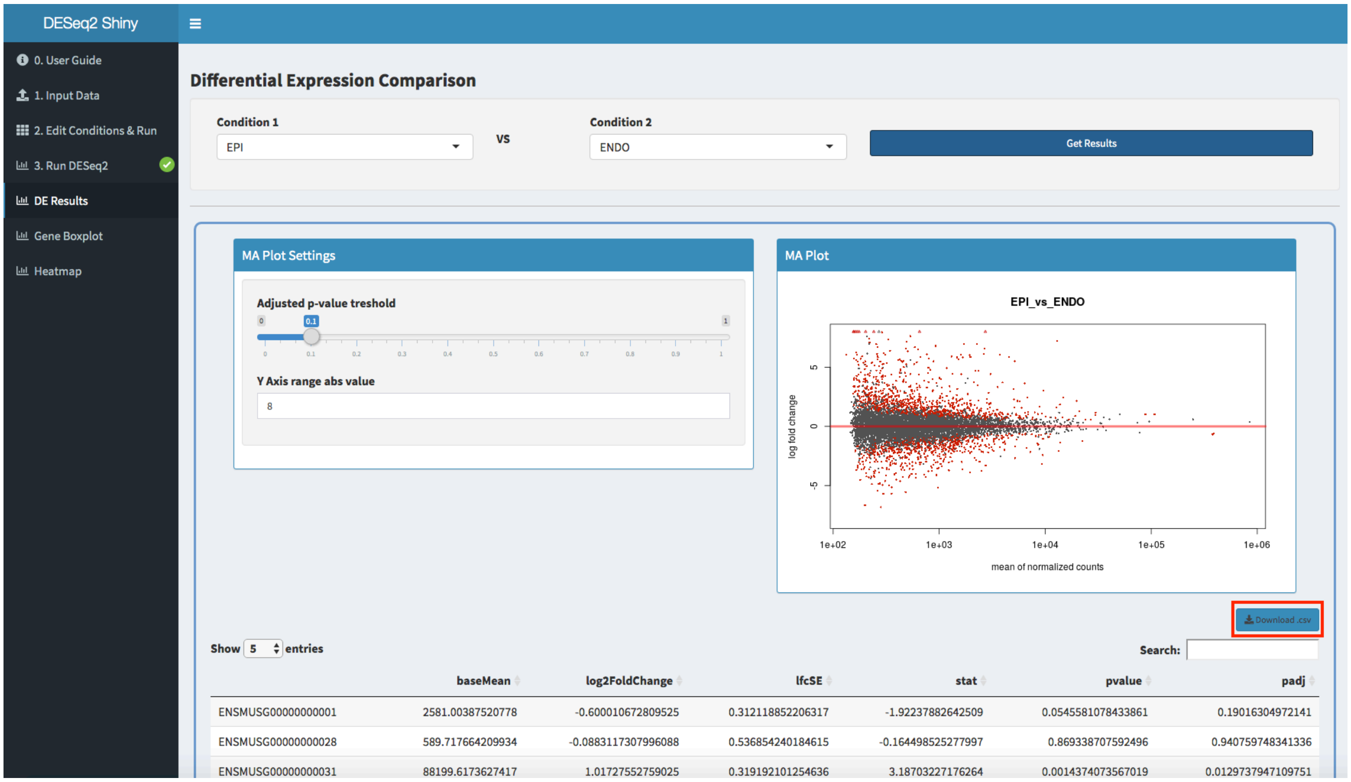

g) There are other visualization plots like gene box plots and clustered heatmap

**Box Plots**:

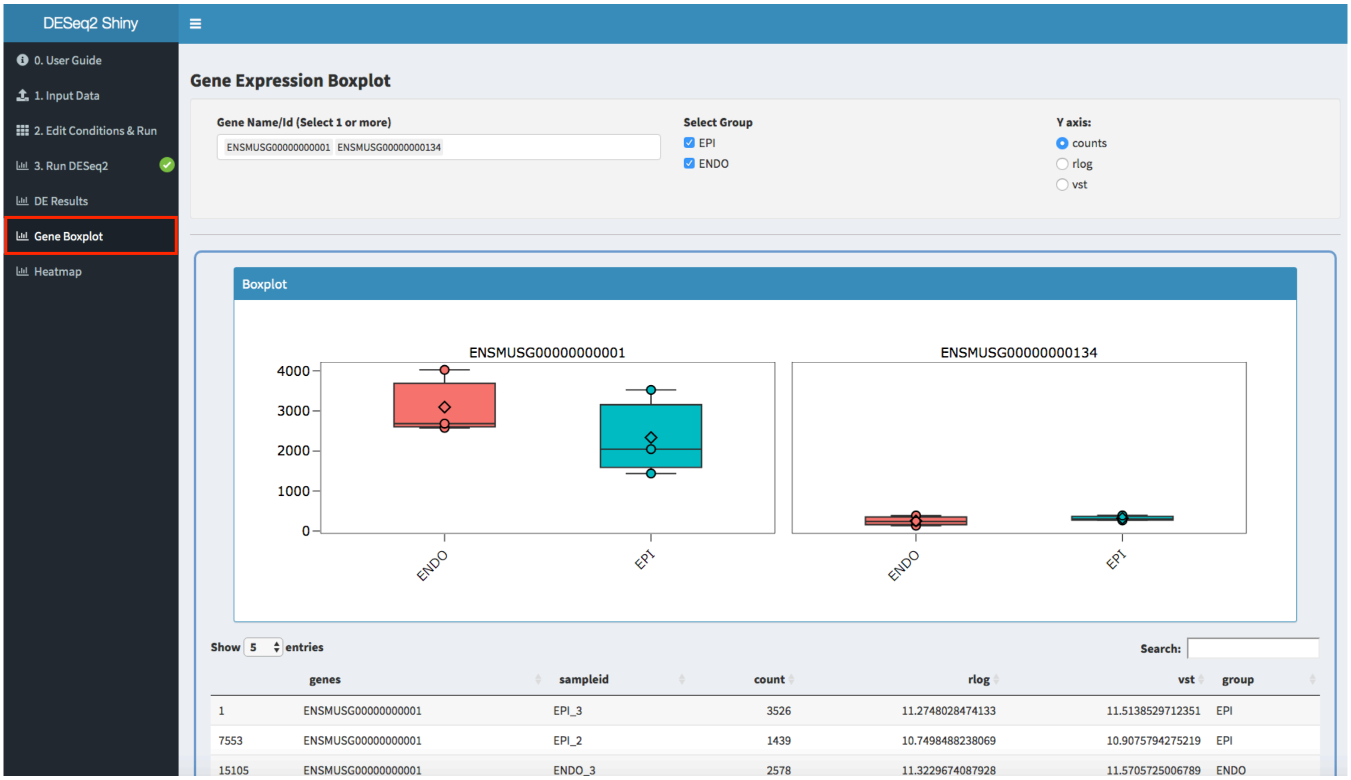


**Clustered Heatmap**:

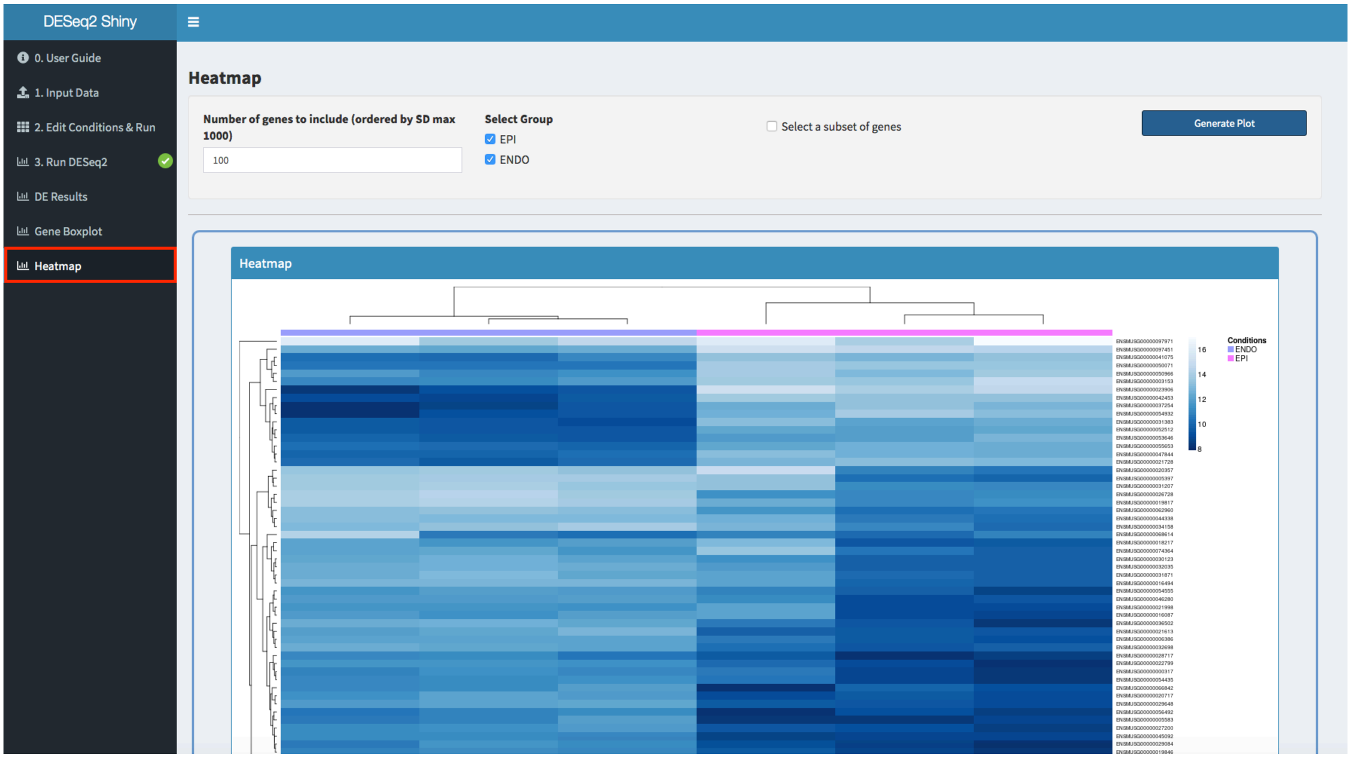

***Step 3: Gene Set Enrichment Analysis (GSEA):***

a) Launch ClusterProfShinyGSEA http://nasqar.abudhabi.nyu.edu/ClusterProfShinyGSEA/

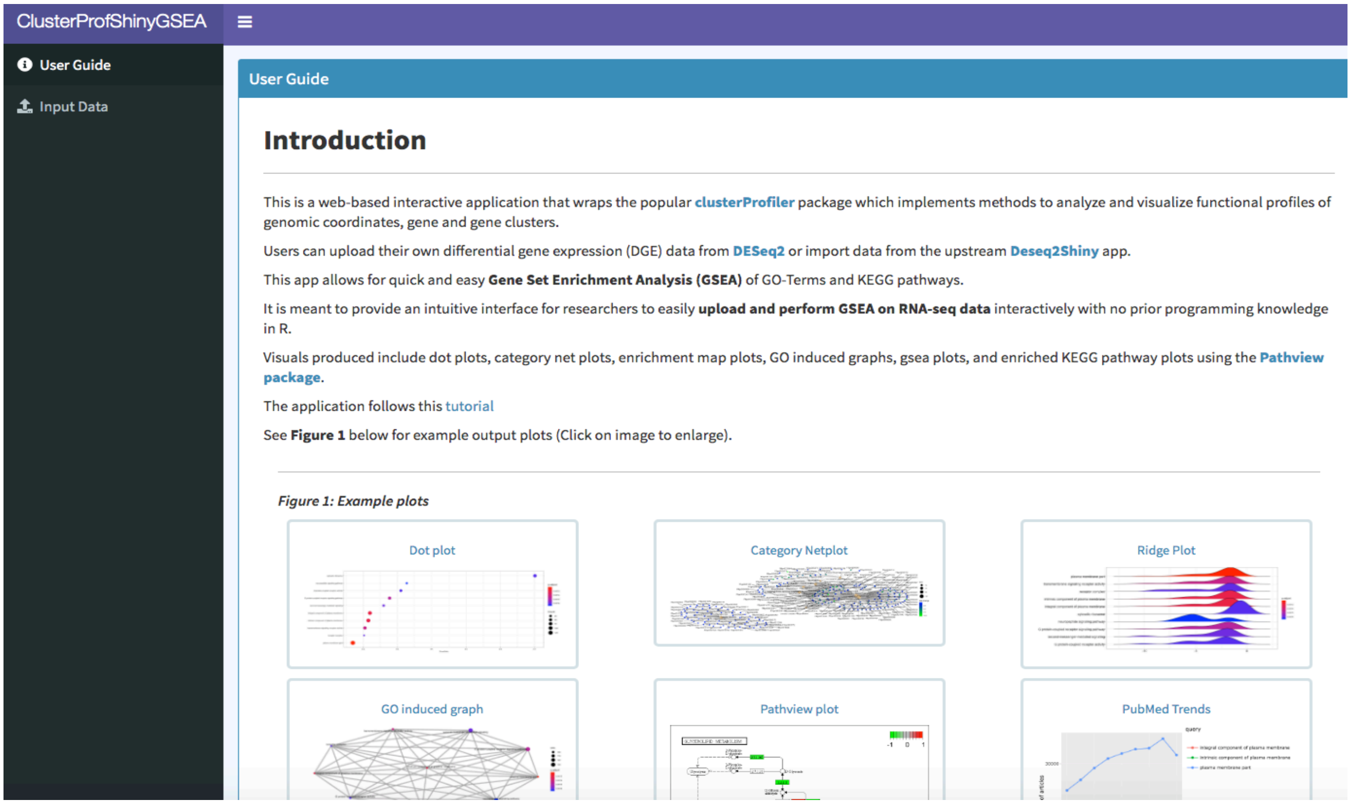


b) Go to "**Input Data**" tab and click "**Browse**"

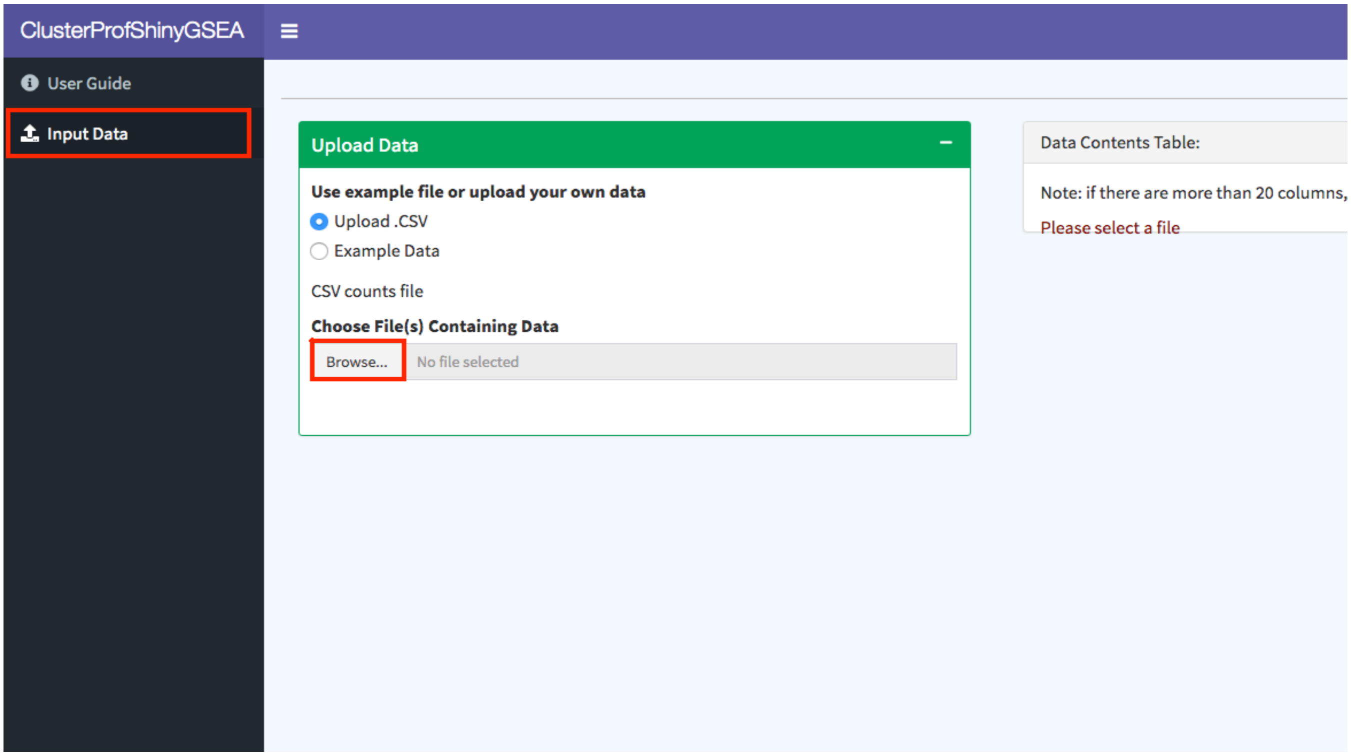

c) Select the DE results file downloaded in the previous section (Section 2, Step f) from Deseq2Shiny (**EPI_vs_ENDO.csv**) to upload.

d) In the tab “Initialize Parameters”, make sure to select the correct column name that corresponds to the LogFC column and click “**Next**”

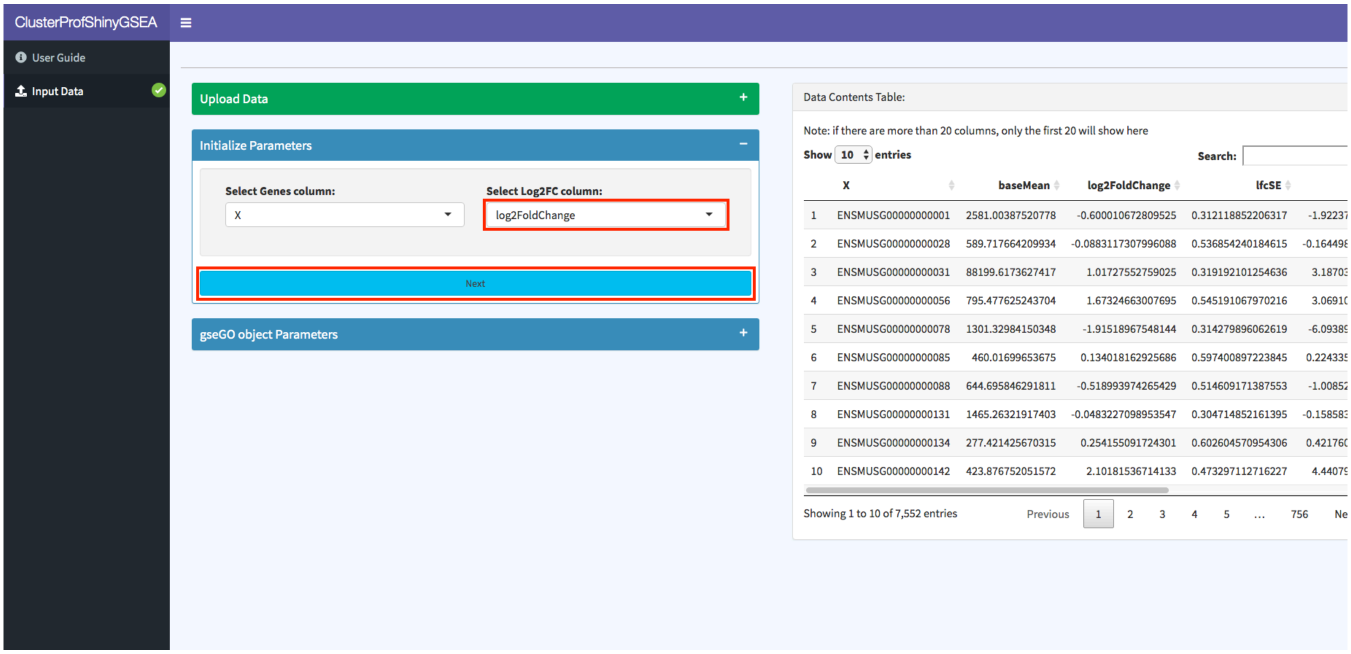


e) Select “**Mouse (org.Mm.eg.db)**” as the organism and click “**Create gseGO Object**” to start the analysis

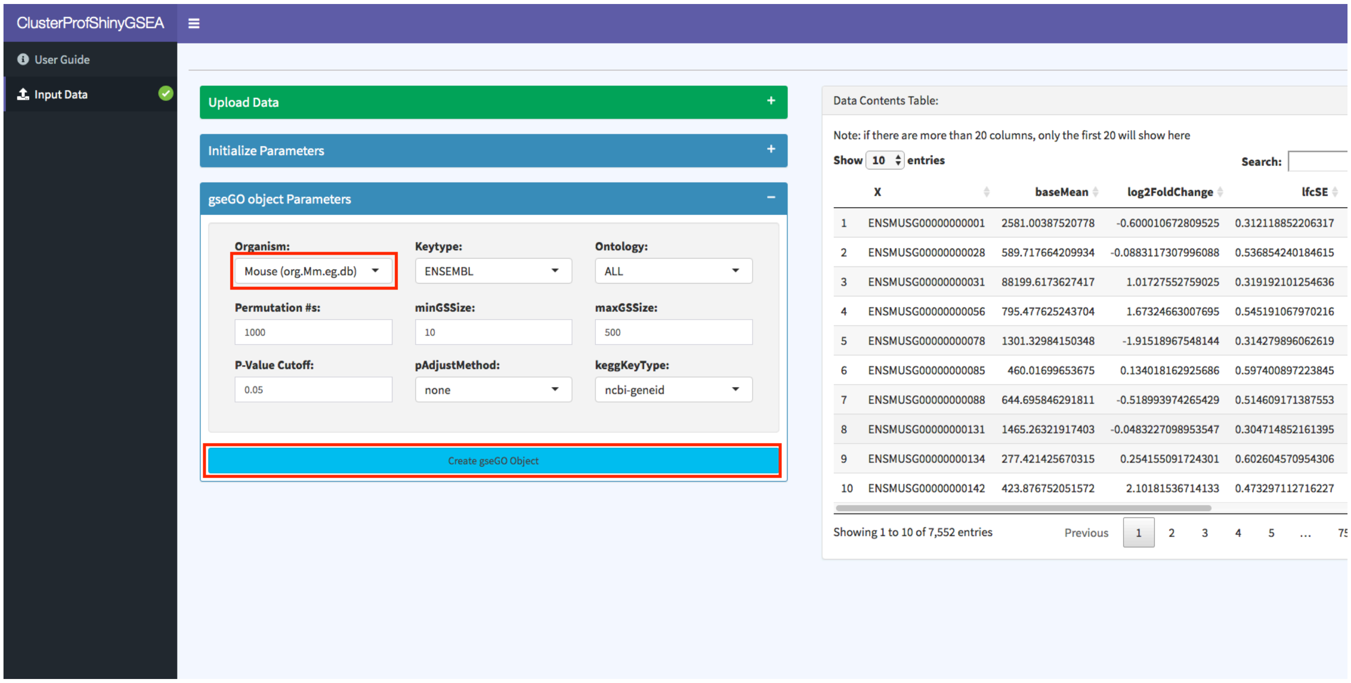

f) Now you can see the gseGO results table. Next go to “**gseKegg Results**” tab to view gseKEGG results table.

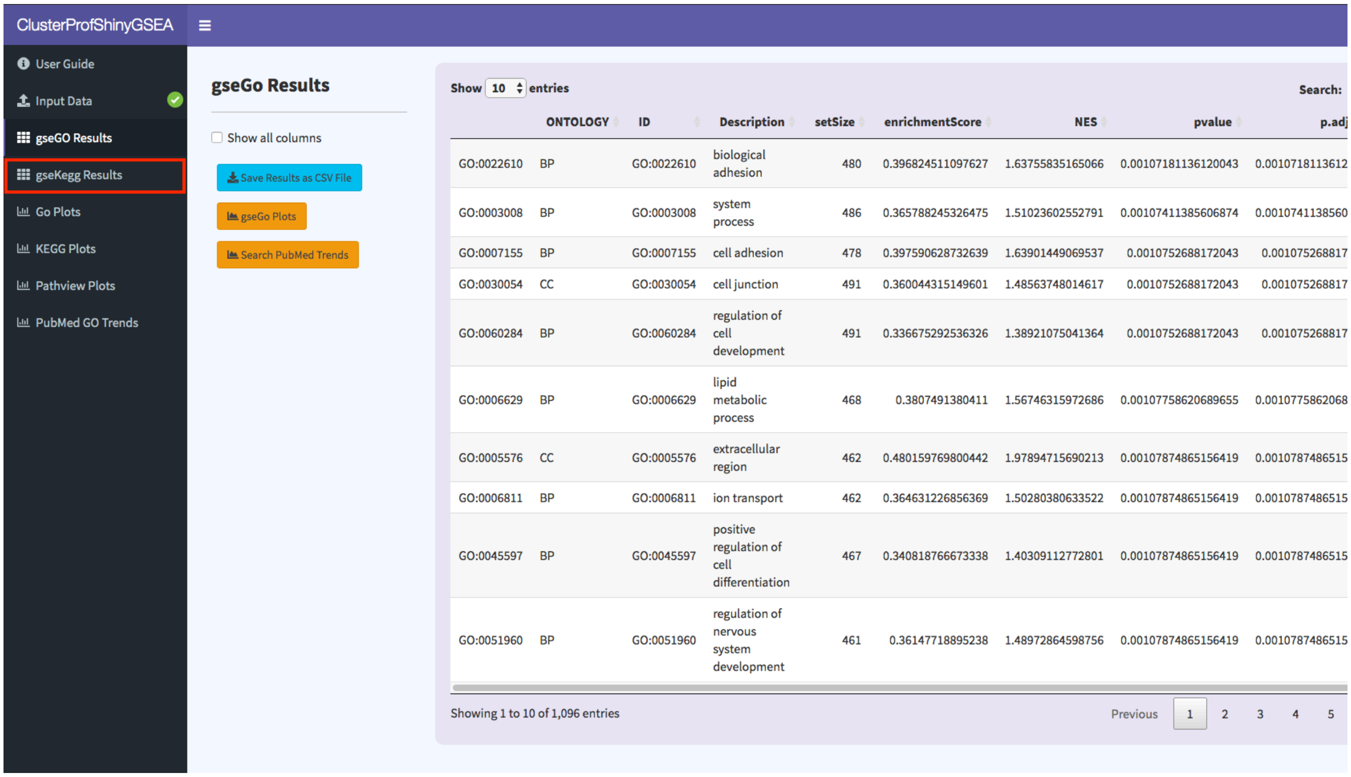

g) Here are the Kegg results with an output that indicates the percentage of genes that were not mapped. Next go to “**GO Plots**” tab

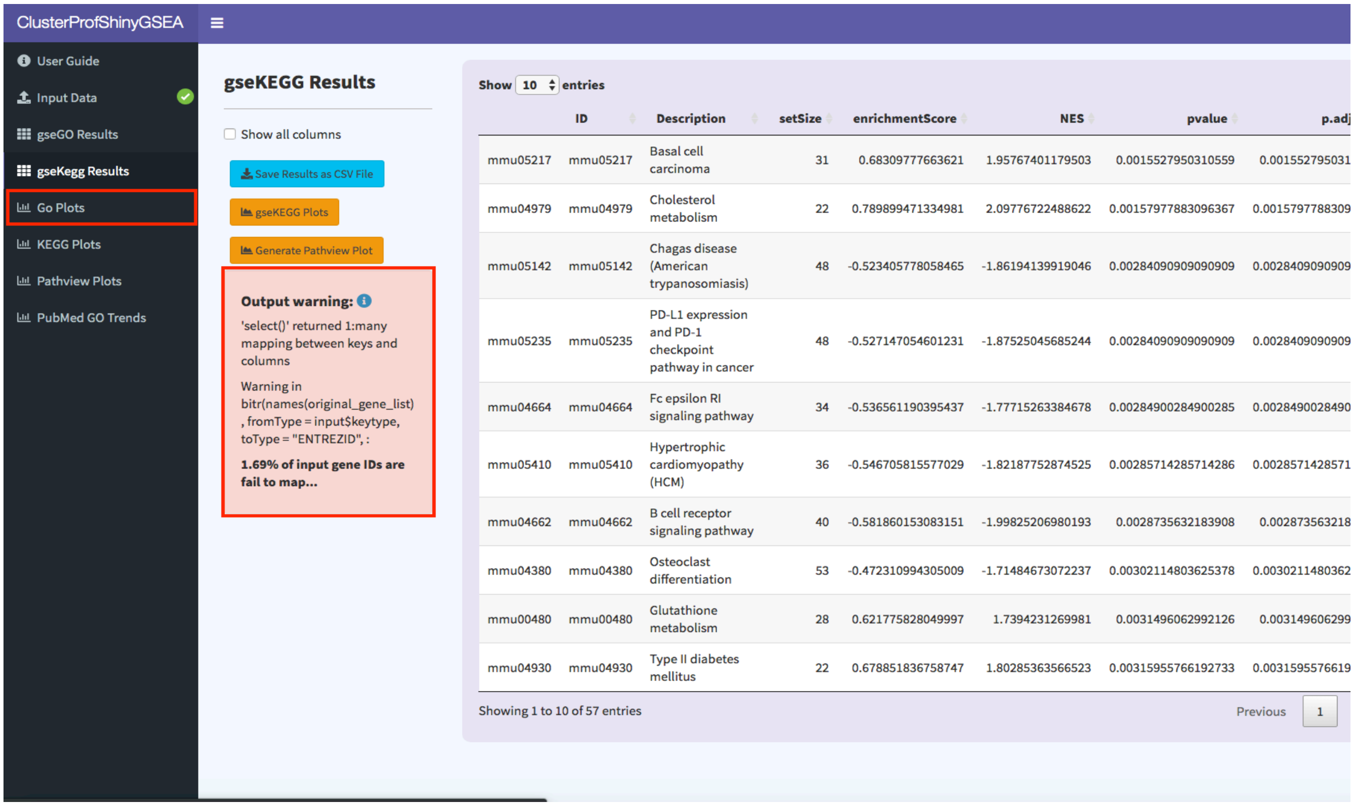


h) There are several plots to explore (Dot plot, Enrichment map)

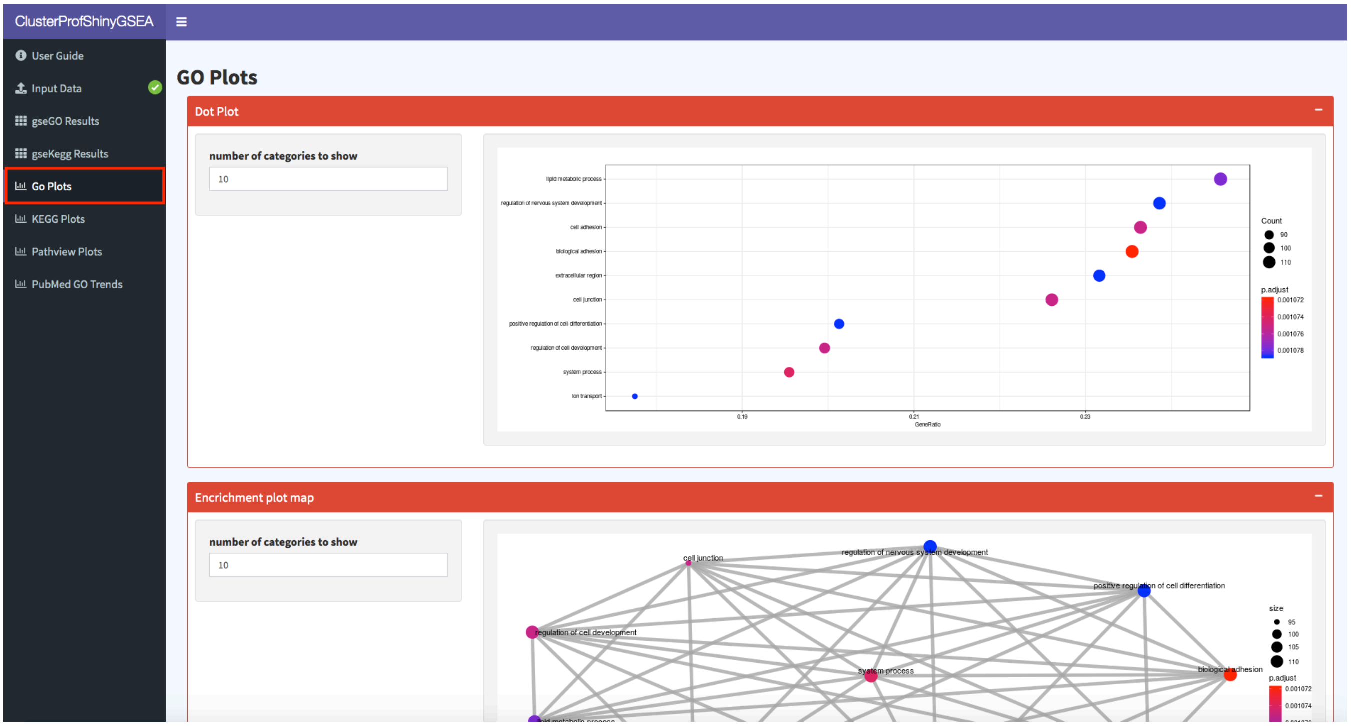

i) Ridge plot and GSEA plot

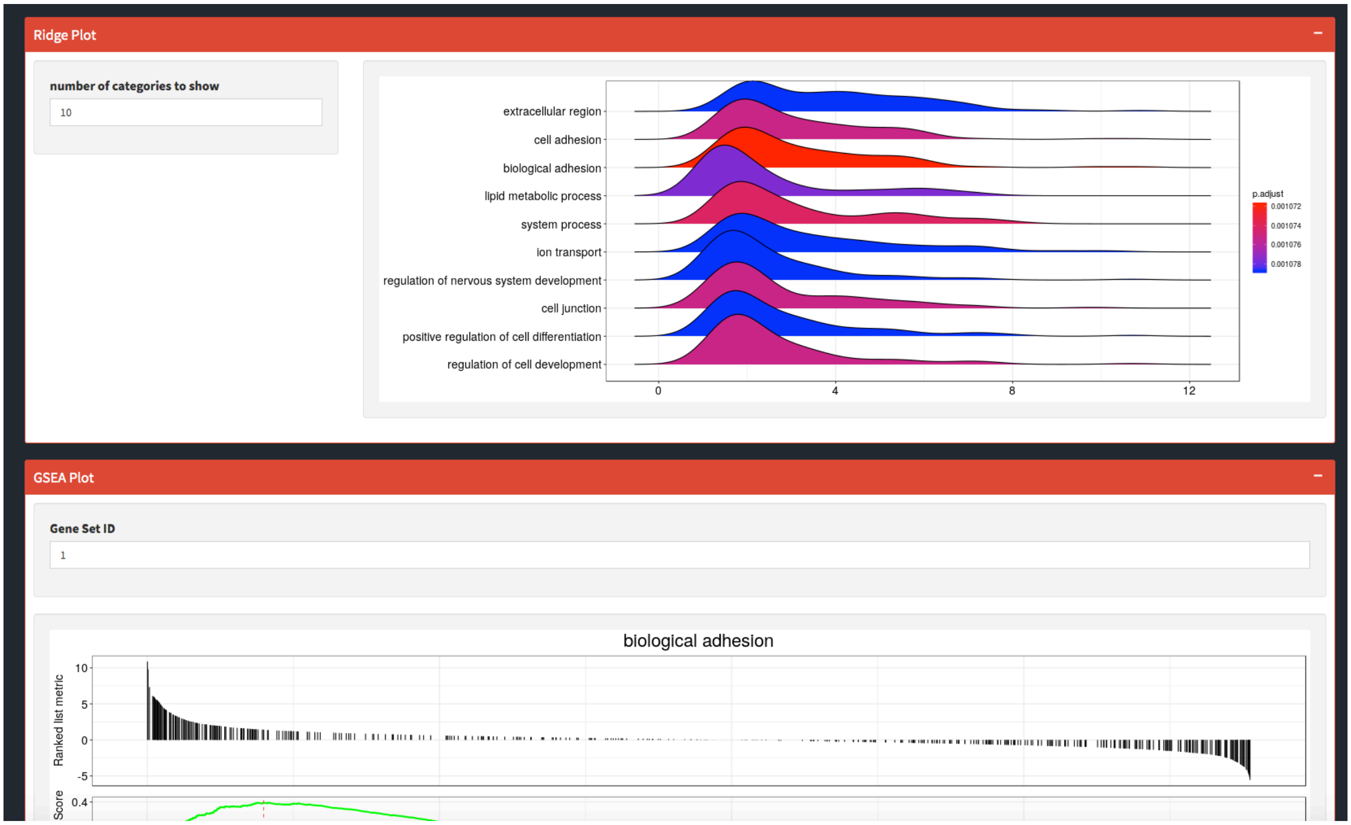


j) Next go to "**Pathview Plots**" tab and select gene "**mmu05217**" and click "**Generate Pathview**"

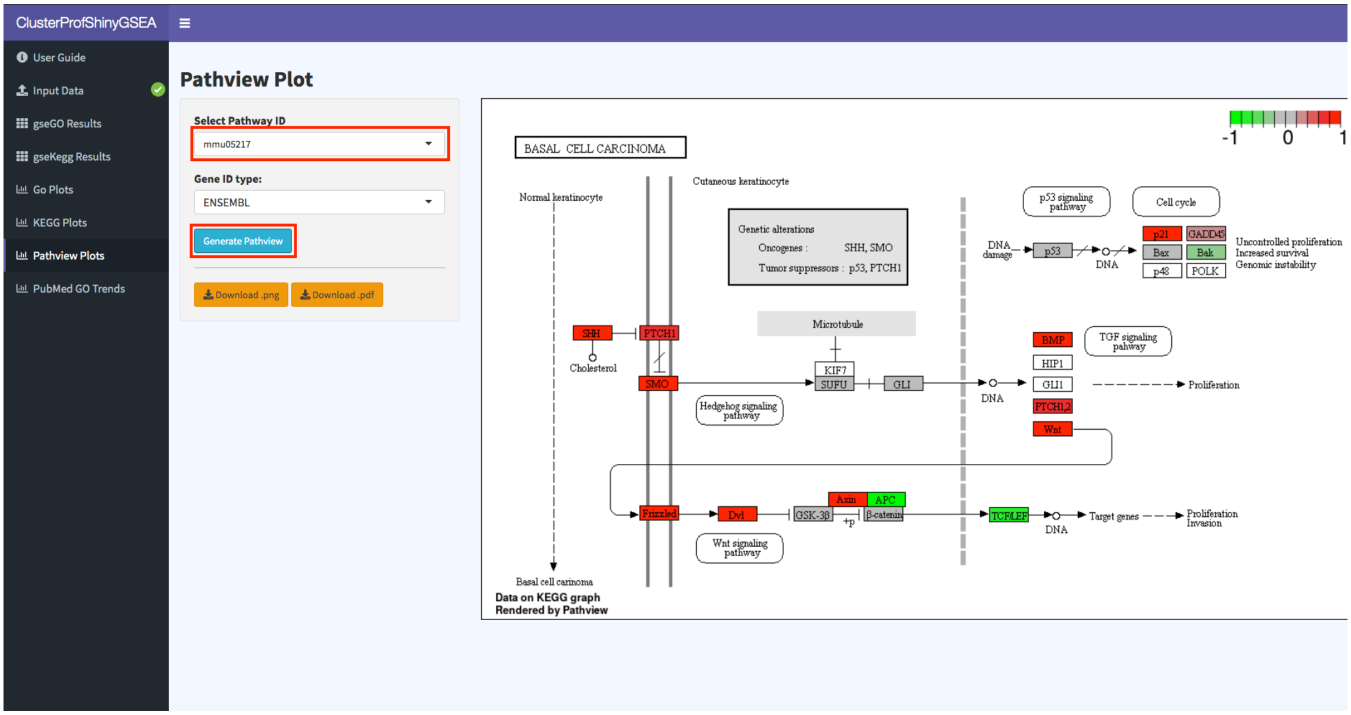

k) Last step is to check pubmed trends for enriched categories. Select a few GO terms and click "**Plot Trends**"

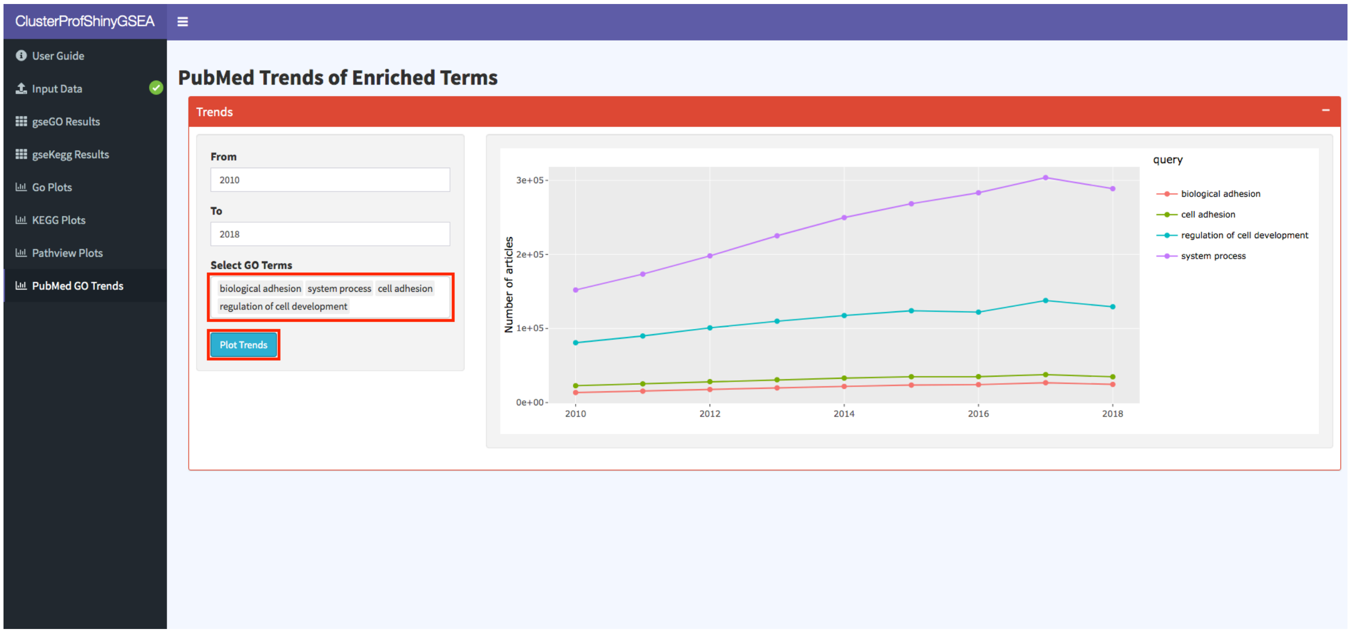

### 4) Example Use Case 2 (scRNA-seq analysis using Seurat):

In this example we will show how you can easily upload your single-cell RNA seq sample data (10X format, using the 10x genomics cellranger pipeline) and perform a guided single-cell data analysis and visualization using the popular Seurat library. You can follow along using the Seurat tutorial (https://satijalab.org/seurat/v3.0/pbmc3k_tutorial.html), which the SeuratV3Wizard mirrors closely.

a) Download and extract data file from https://s3-us-west-2.amazonaws.com/10x.files/samples/cell/pbmc3k/pbmc3k_filtered_gene_bc_matrices.tar.gz
b) Launch SeuratV3Wizard (http://nasqar.abudhabi.nyu.edu/SeuratV3Wizard/)

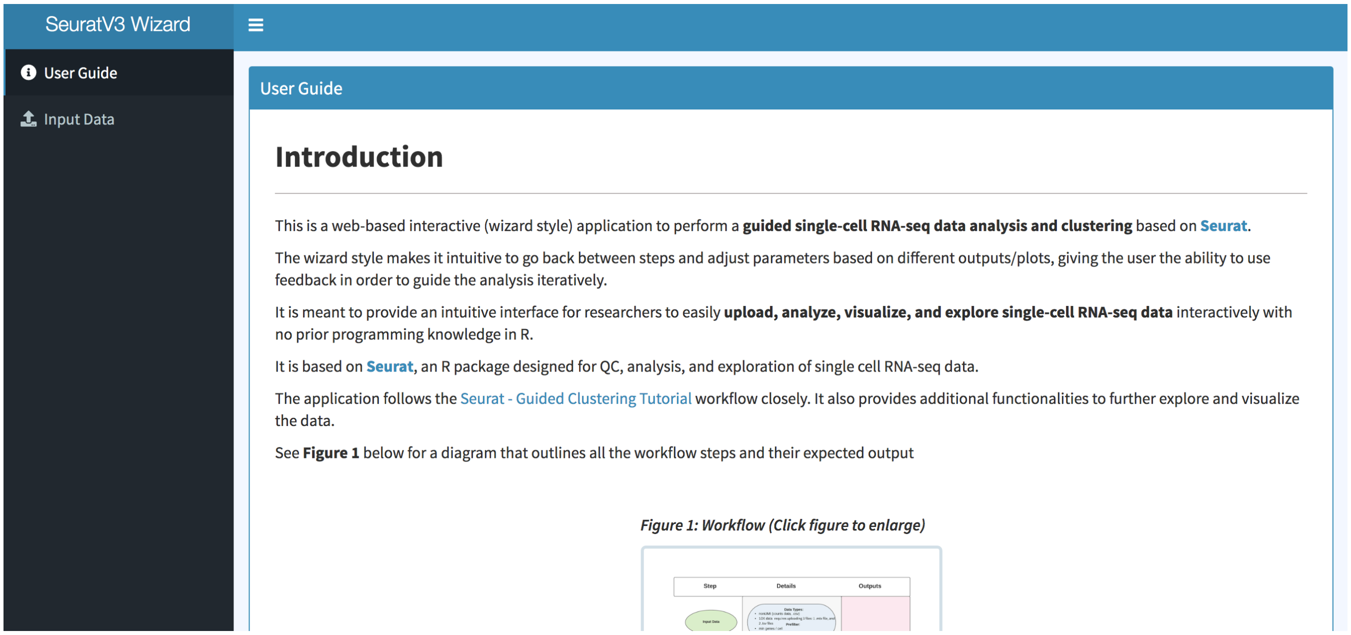

c) Go to “**Input Data**” tab and select “**Upload Data (10X)**”

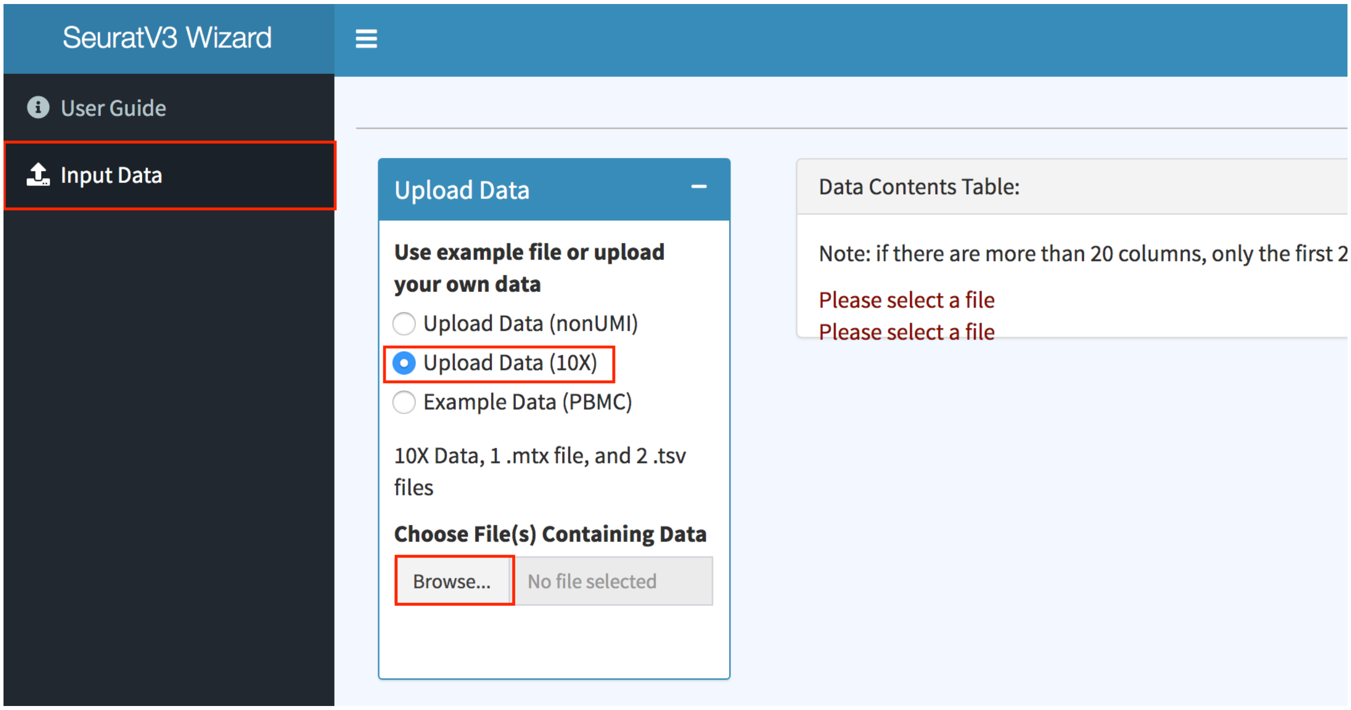


d) Browse for the extracted data files downloaded earlier. Select all 3 files to upload

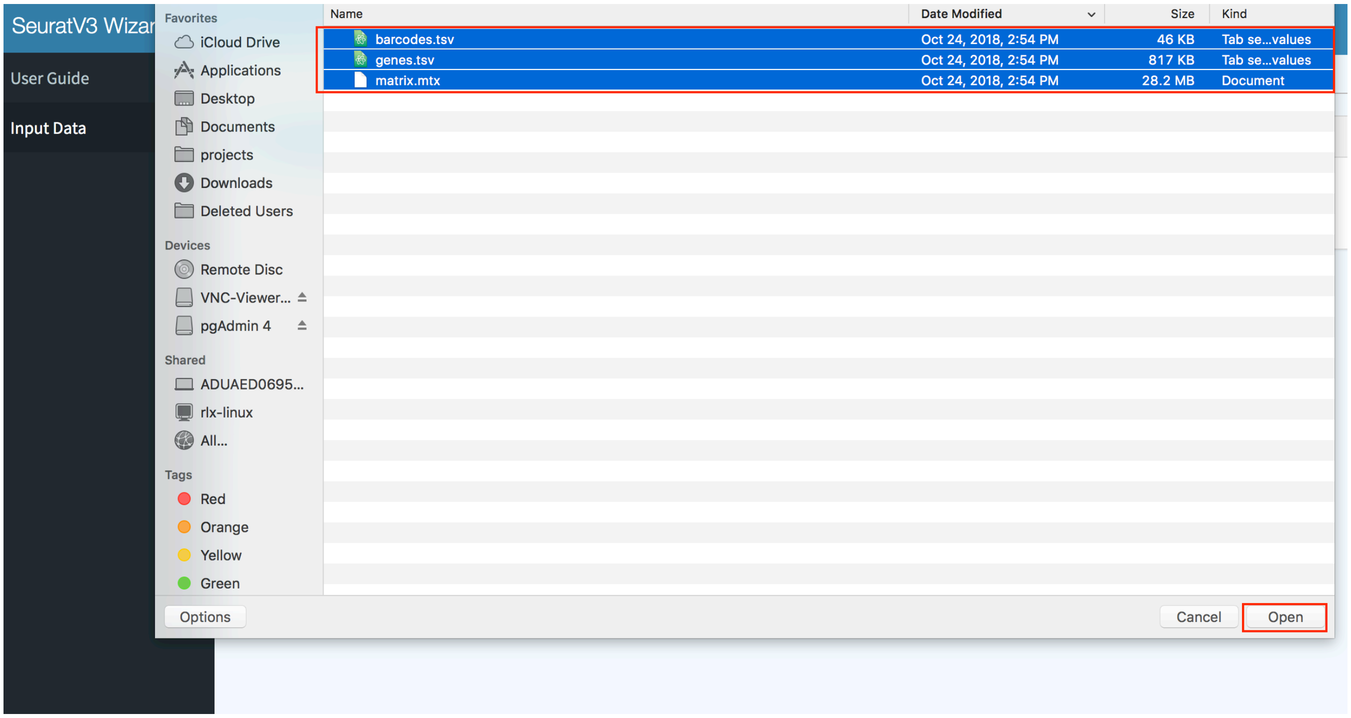

e) Click “**Next Step: QC & Filter Cells**”

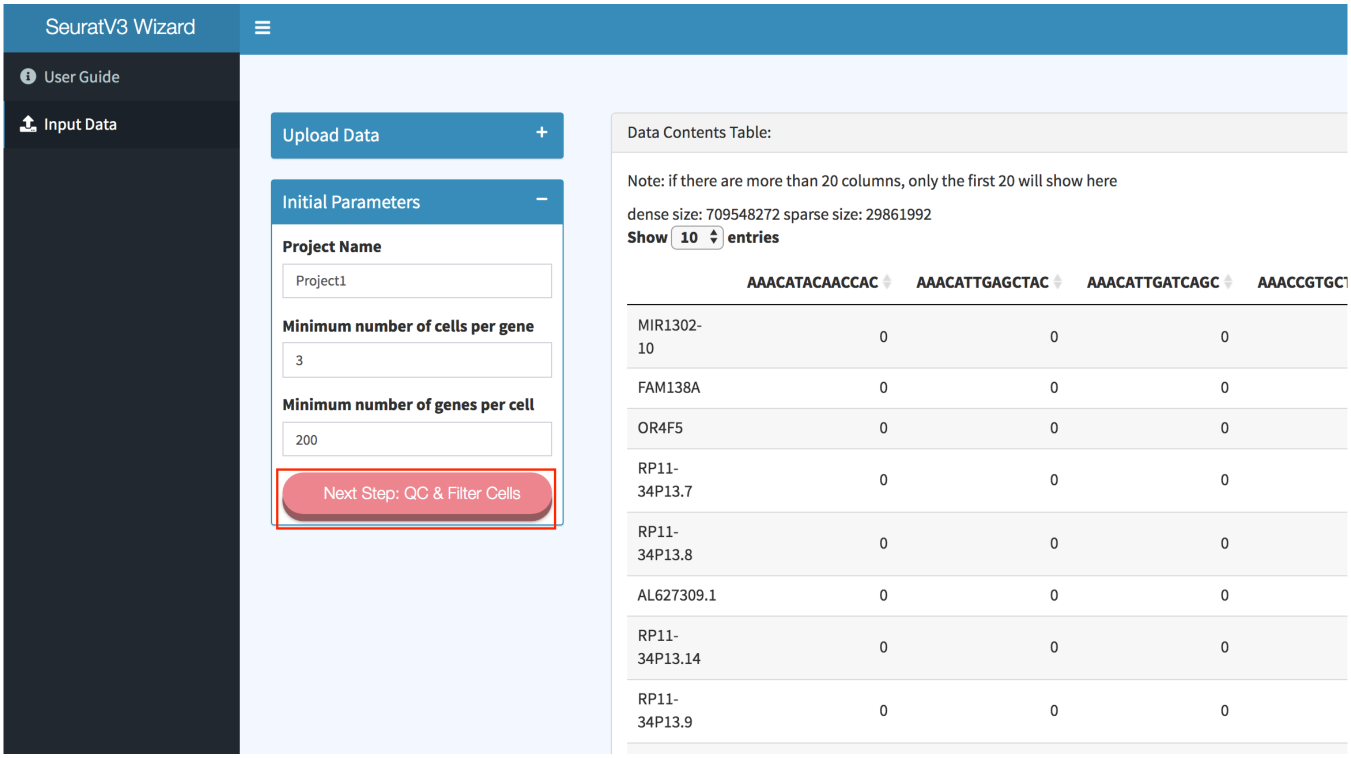


f) Add a regular expression for mitochondrial genes by adding the **regex** and **label** as seen below, and then click “**Add Filter**”. You can use the “**Test Regex”** tab in order to verify that the regular expression the user supplies works as expected.

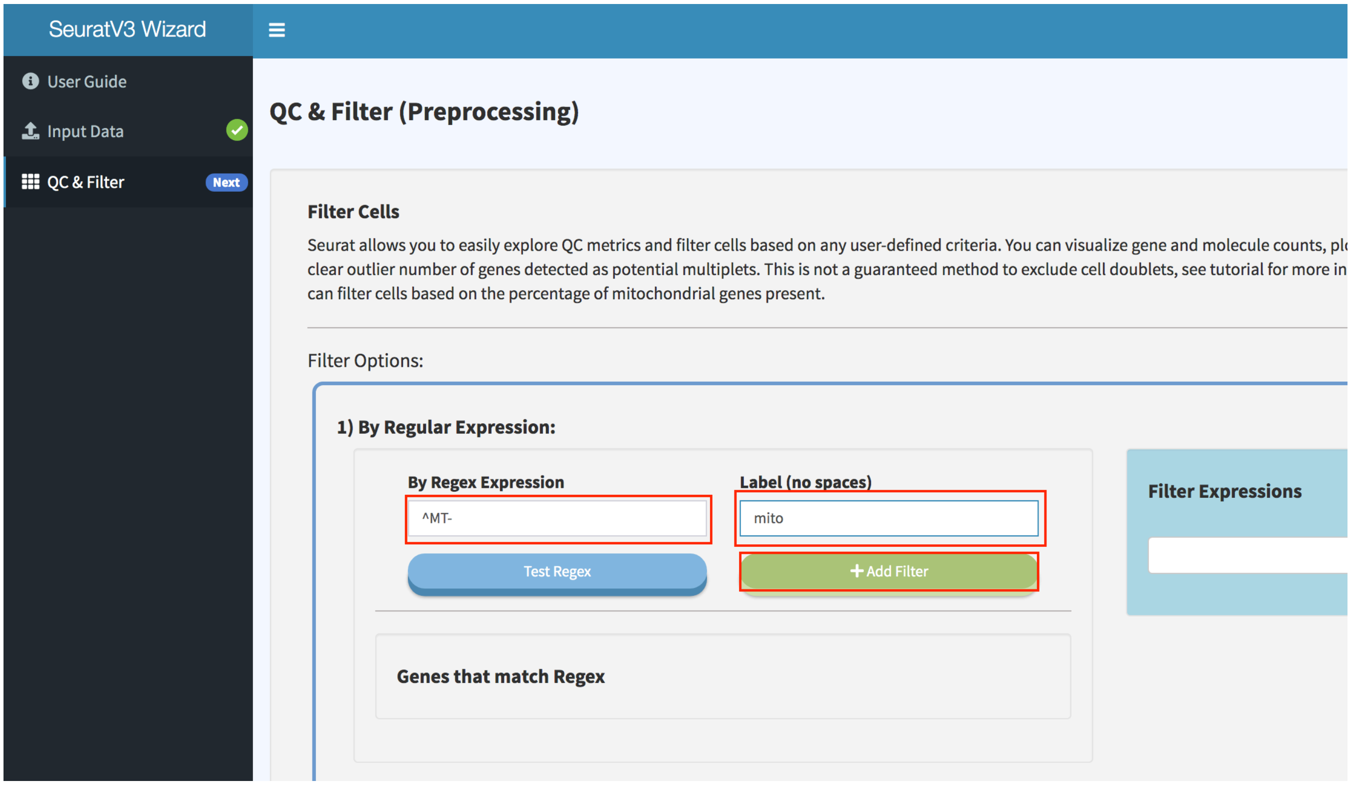

g) Scroll down and click “**Submit Data**”

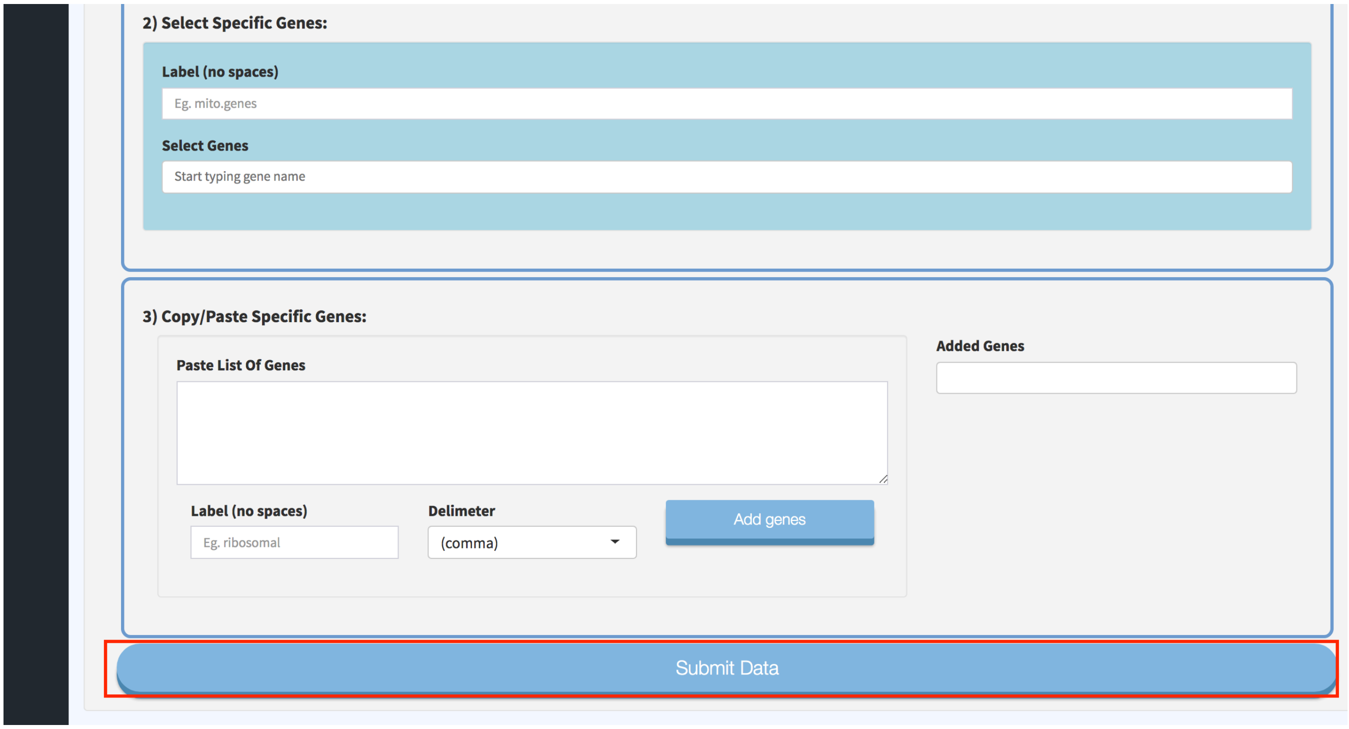


h) Select the low and high thresholds to filter out cells. Click “**Filter Cells (within thresholds)**”. This is an interactive filter that will display the effects of applying different filtering thresholds on the data.

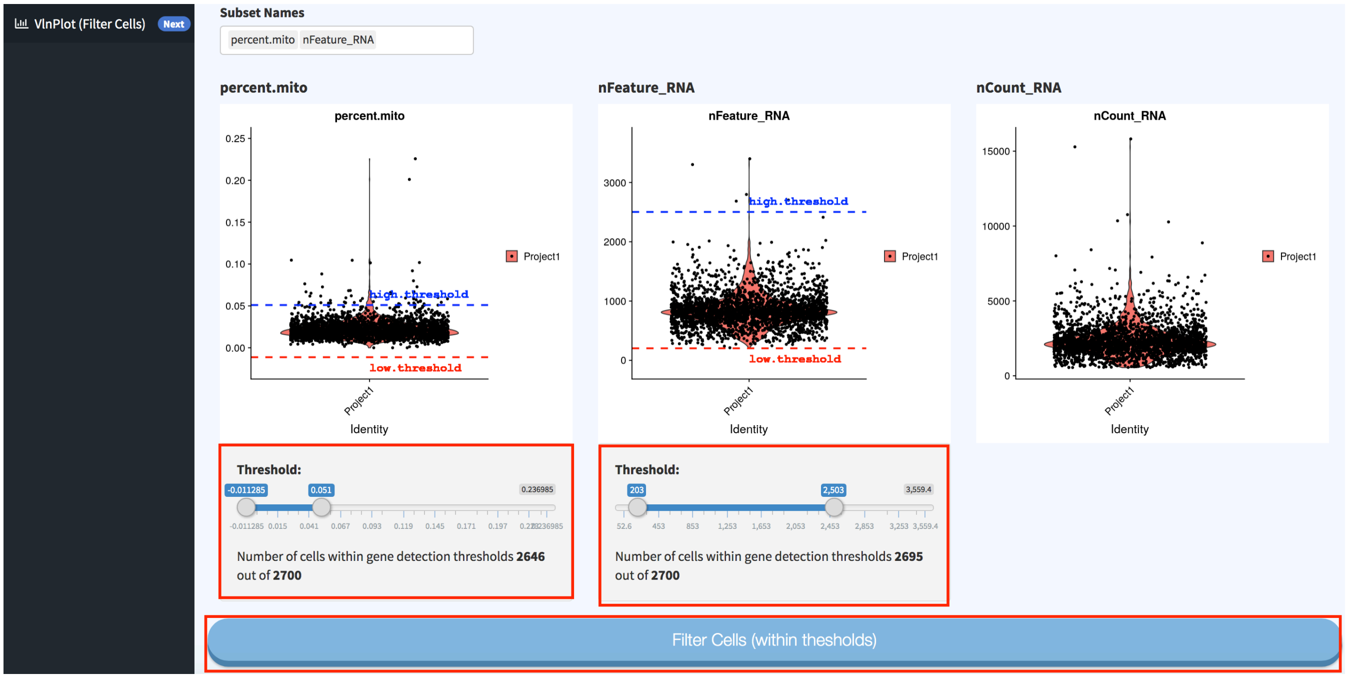

i) There are now two options to proceed within the analysis. We will choose the default one so we can mirror the tutorial. The second option is the SCTransform method (https://rawgit.com/ChristophH/sctransform/master/inst/doc/seurat.html )

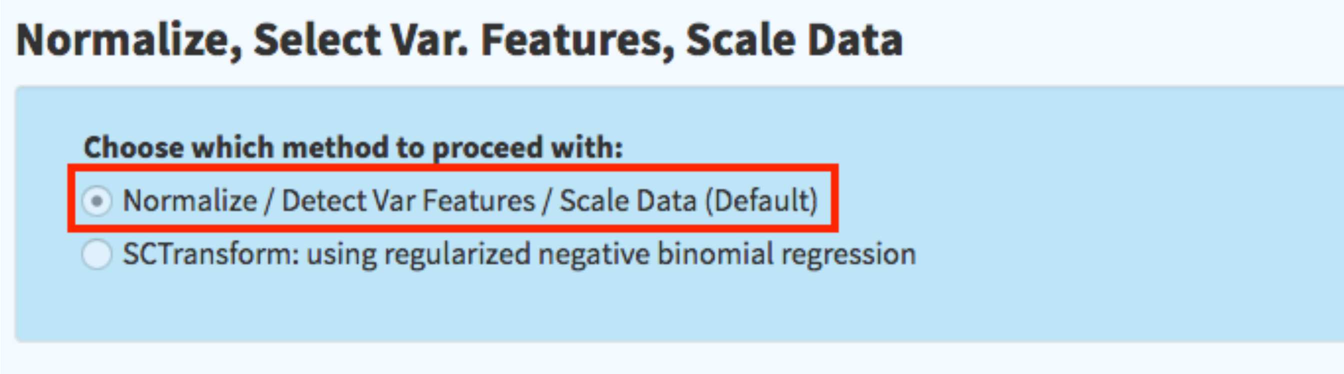


j) Proceed by clicking “**Normalize / Find Var. Feaatures / Scala Data**”

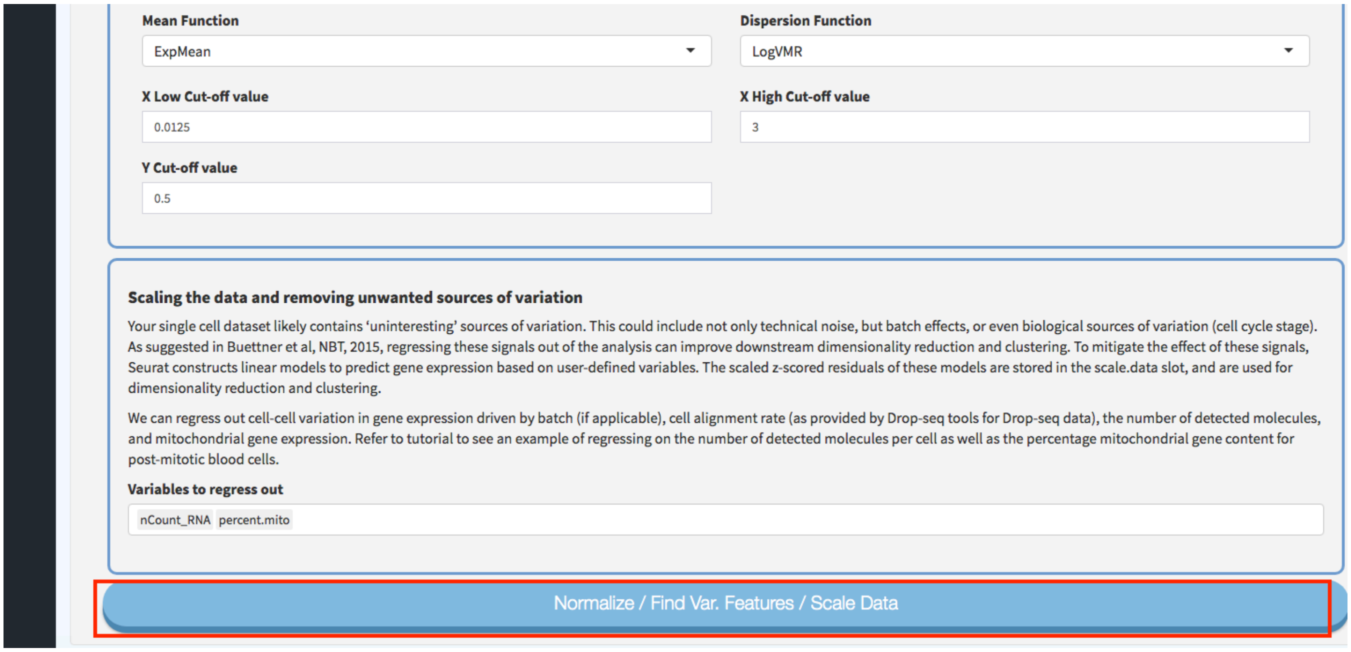

k) Now you can follow the default settings in the wizard to go through the steps until "**Elbow/Jackstraw**". Click "**Show Elbow Plot**" and click "**Next Step: Cluster Cells**"

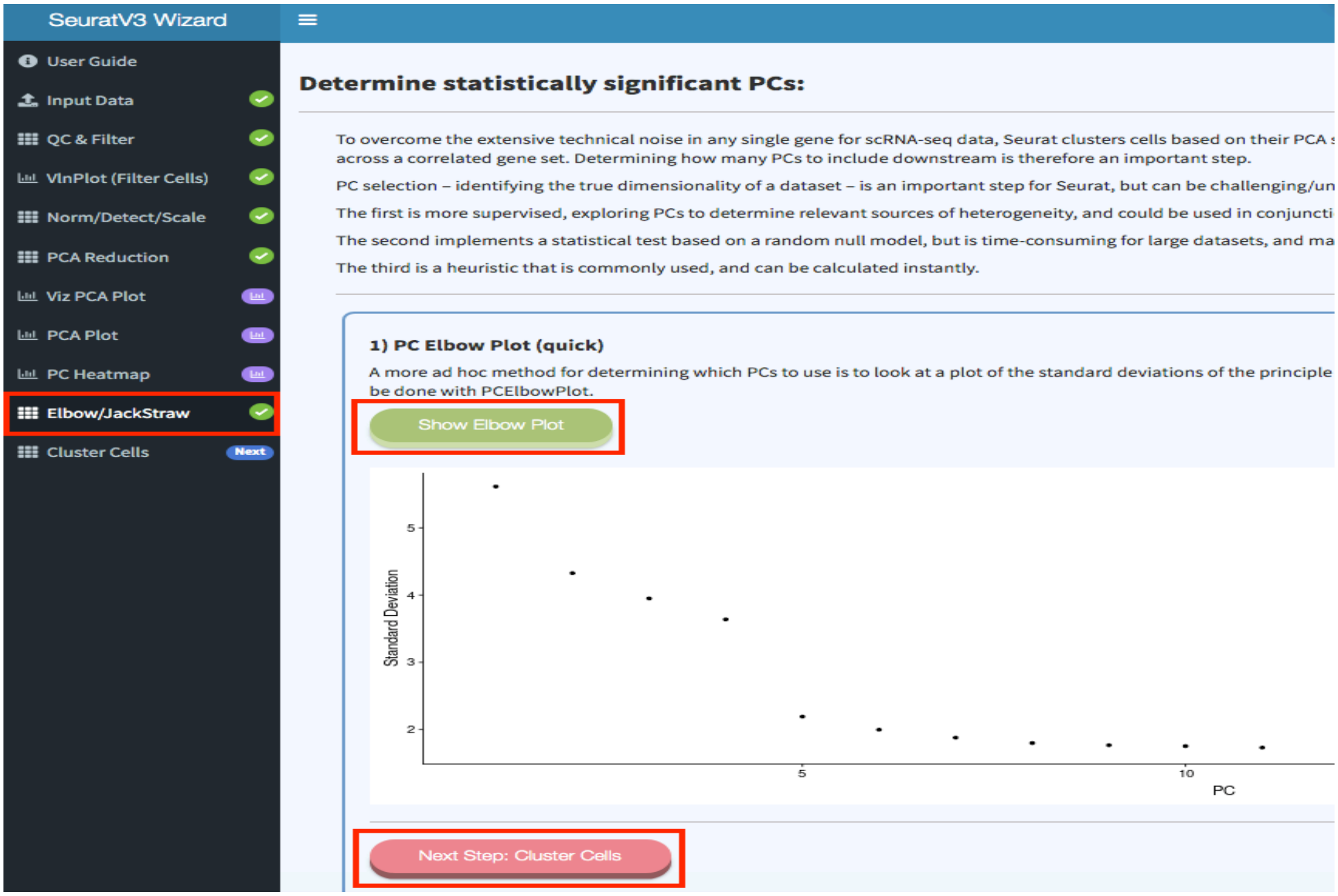

l) Next run "**Cluster Cells**" and go to "**Non-linear Reduction**" tab and "**Run TSNE Reduction**"

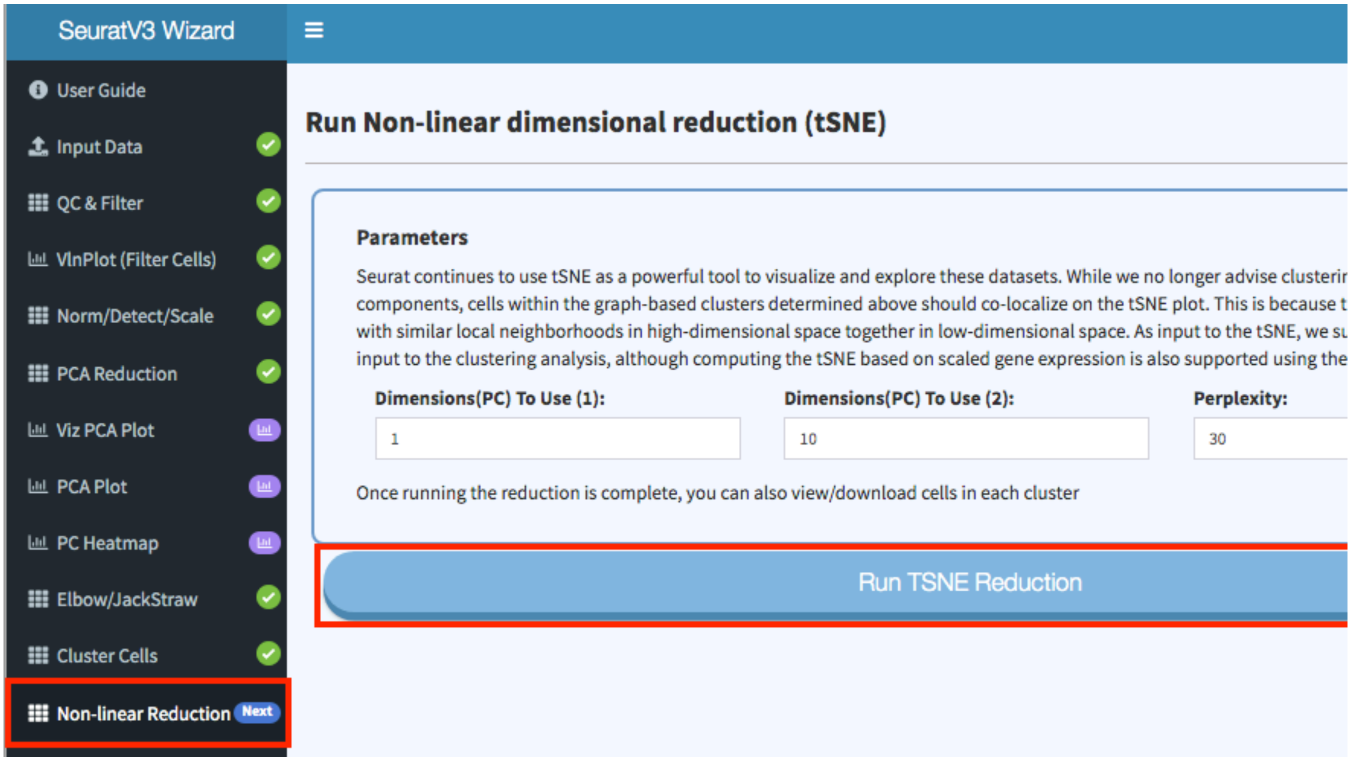


m) Once tSNE step is done, we can now go to the next step "**Next Step: Find Cluster Markers**" and click "**Find Cluster Markers**"

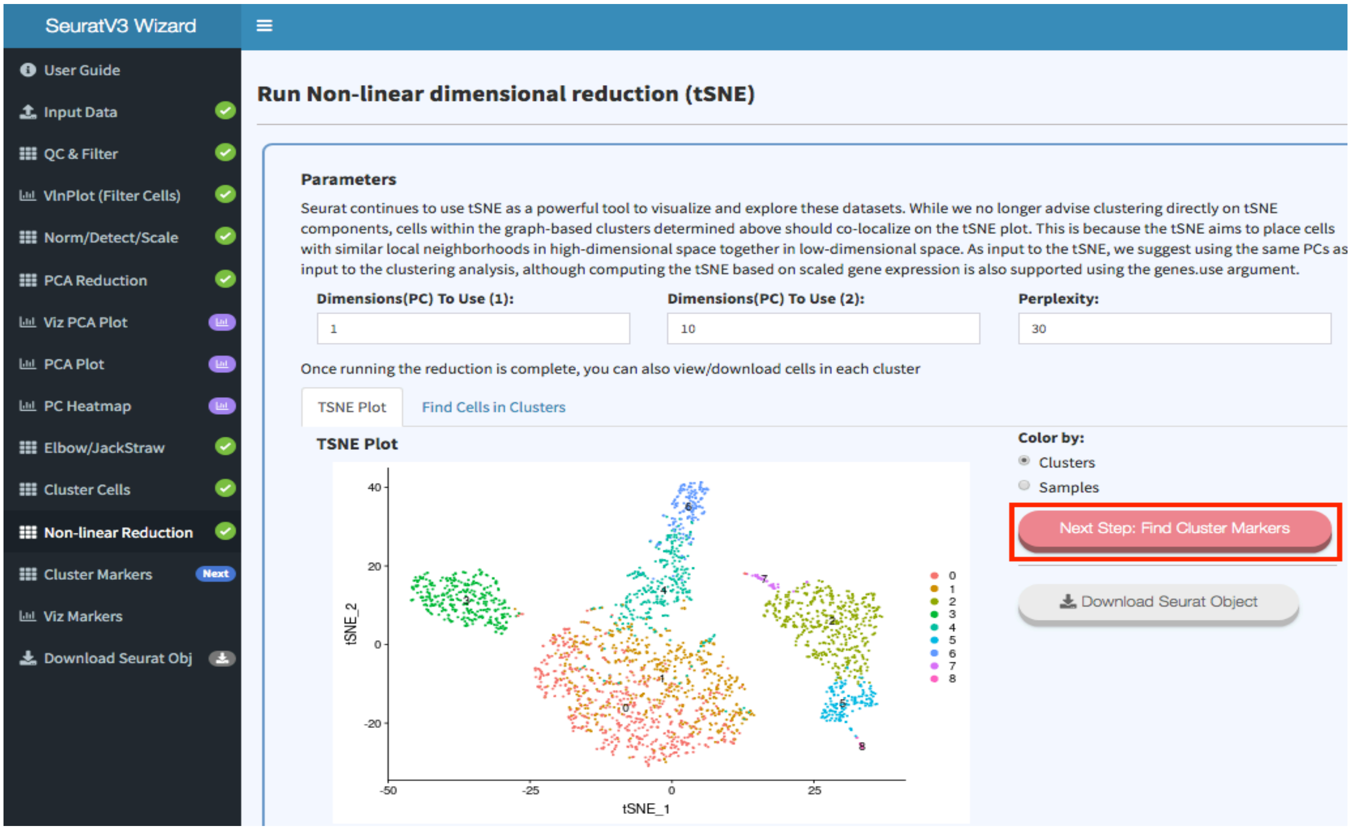

n) Now you can see the markers table (which is downloadable). There is also an option to explore the clusters and markers visually using UCSC Cellbrowser (https://github.com/maximilianh/cellBrowser ). Click "**Generate Cell browser data**". Once done you can launch the cellbrowser by clicking on "**Launch Cellbrowser**".

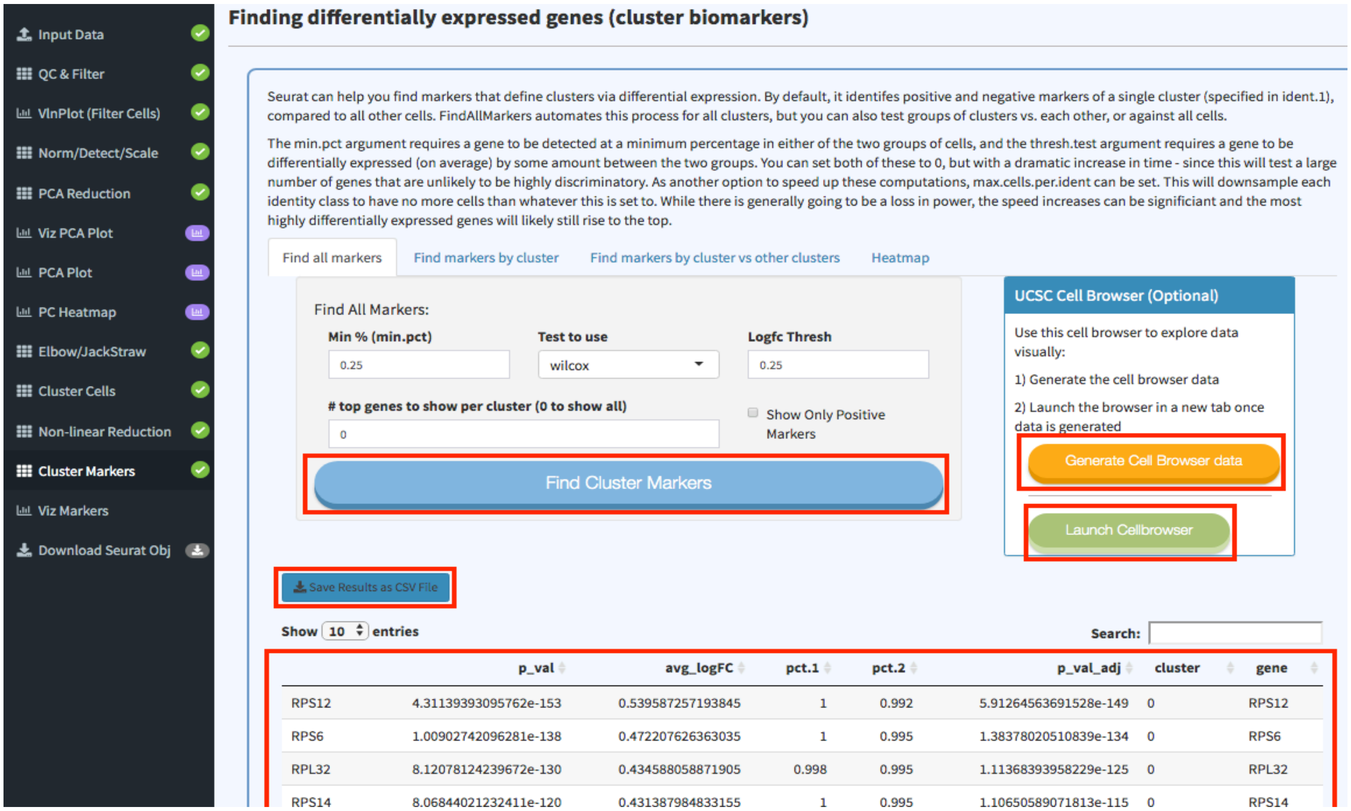

o) A tab is opened with UCSC cellbrowser

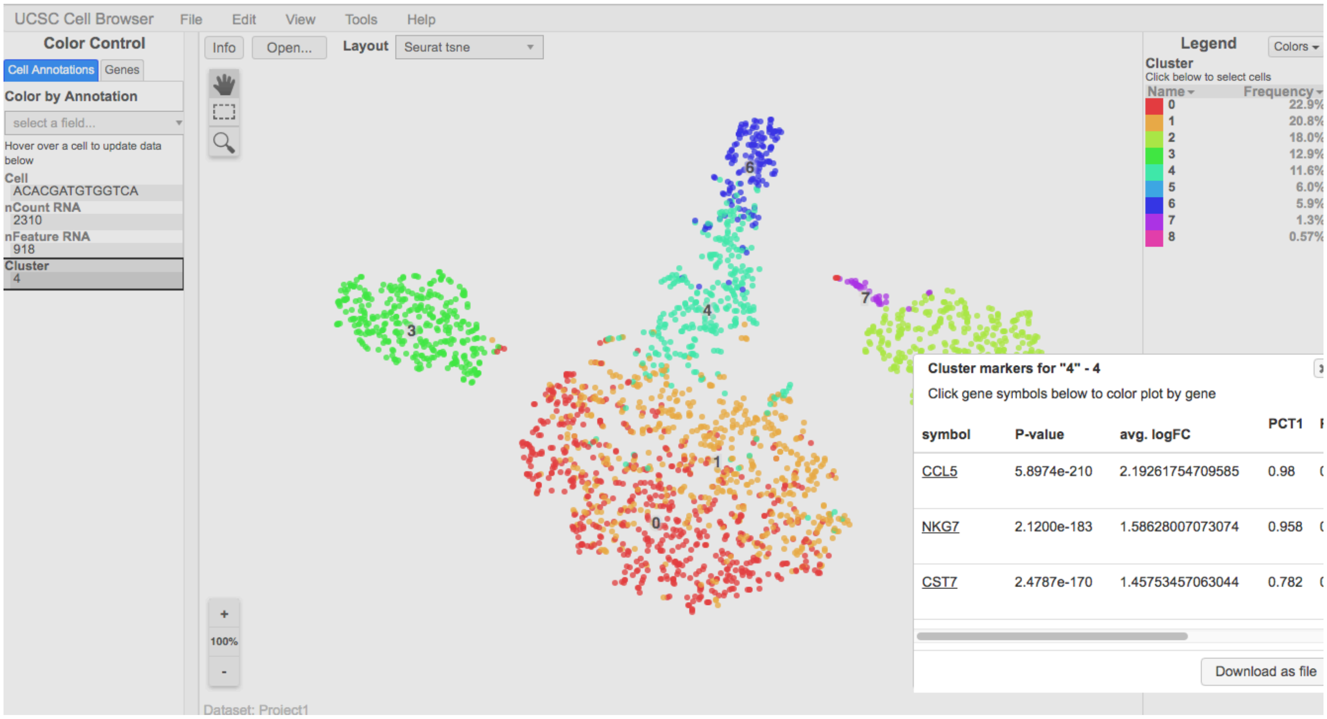


p) Last step would be to download the Seurat object for reproducibility and further analysis in R. Go to the last tab "**Download Seurat Obj**". Click "**Generate Seurat Obj**", once done click "**Download Seurat Obj**" when it appears.
You can also download the R script used for generating this analysis but clicking **"Generate Seurat Script"**.

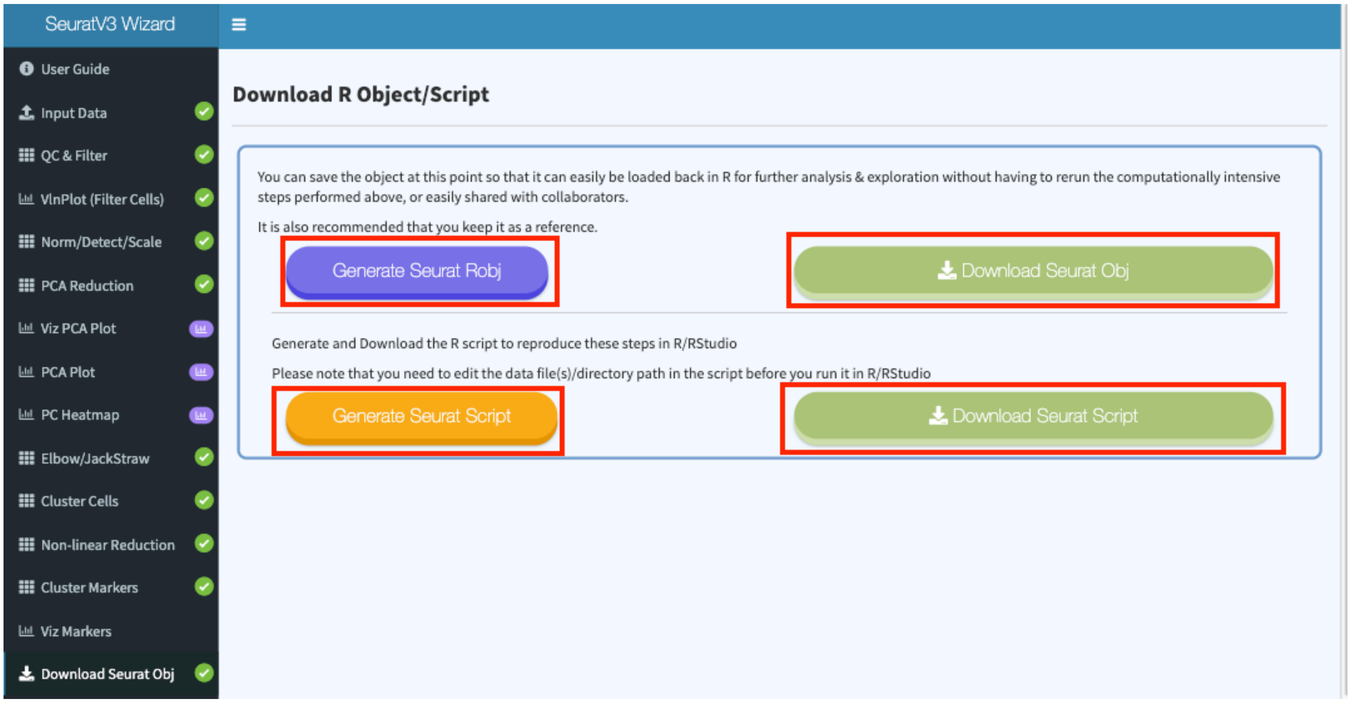

### References


Merkel, D. (2014). Docker: Lightweight linux containers for consistent development and deployment. *Linux J.*, 2014(239).

Soppelsa, F. and Kaewkasi, C. (2017). *Native Docker Clustering with Swarm*. Packt Publishing.

R Core Team (2017). *R: A Language and Environment for Statistical Computing*. R Foundation for Statistical Computing, https://www.R-project.org/.

Chang, W., Cheng, J., Allaire, J., Xie, Y., and McPherson, J. (2018). *shiny: Web Application Framework for R*. R package version 1.1.0.

Love, M. I., Huber, W., and Anders, S. (2014). Moderated estimation of fold change and dispersion for rna-seq data with deseq2. *Genome Biology*, 15, 550.

Butler, A., Hoffman, P., Smibert, P., Papalexi, E., and Satija, R. (2018). Integrating single-cell transcriptomic data across different conditions, technologies, and species. *Nature Biotechnology*, 36, 411.

Pyl, P. T., Anders, S., and Huber, W. (2014). HTSeq—a Python framework to work with high-throughput sequencing data. *Bioinformatics*, 31(2), 166–169.

Smyth, G. K., Shi, W., and Liao, Y. (2013). featureCounts: an efficient general purpose program for assigning sequence reads to genomic features. *Bioinformatics*, 30(7), 923–930.

Sklenar, J., Nelson, J. W., Minnier, J., and Barnes, A. P. (2016). The START App: a web-based RNAseq analysis and visualization resource. *Bioinformatics*, 33(3), 447–449.

Stuart, T. and Satija, R. (2019). Integrative single-cell analysis. *Nature Reviews Genetics*, 20(5), 257–272.

Patel, M. V. (2018). iS-CellR: a user-friendly tool for analyzing and visualizing single-cell RNA sequencing data. *Bioinformatics*, 34(24), 4305–4306.

Hafemeister, C. and Satija, R. (2019). Normalization and variance stabilization of single-cell rna-seq data using regularized negative binomial regression. *bioRxiv*.

Yu, G., Wang, L.-G., Han, Y., and He, Q.-Y. (2012). clusterprofiler: an r package for comparing biological themes among gene clusters. *OMICS: A Journal of Integrative Biology*, 16(5), 284–287.

Luo, W. and Brouwer, C. (2013). Pathview: an R/Bioconductor package for pathway- Li, Y. and

Andrade, J. (2017). Deapp: an interactive web interface for differential expression analysis of next generation sequence data. *Source Code for Biology and Medicine*, 12(1), 2._based data integration and visualization. *Bioinformatics*, 29(14), 1830–1831.

Quereda, J. J., Dussurget, O., Nahori, M.-A., Ghozlane, A., Volant, S., Dillies, M.-A., Regnault, B., Kennedy, S., Mondot, S., Villoing, B., Cossart, P., and Pizarro-Cerda, J. (2016). Bacteriocin from epidemic listeria strains alters the host intestinal microbiota to favor infection. *Proceedings of the National Academy of Sciences*, 113(20), 5706–5711.

Choi, E. , Kraus, M. R., Lemaire, L. A., Yoshimoto, M. , Vemula, S. , Potter, L. A., Manduchi, E. , Stoeckert, C. J., Grapin_Botton, A. and Magnuson, M. A. (2012), Dual Lineage_Specific Expression of Sox17 During Mouse Embryogenesis. STEM CELLS, 30: 2297-2308.

[dataset] (2016). Transcription profiling by high throughput sequencing of Sox17.Epi and Endo cells from mouse embryos, atlas-experiments, V1. http://www.ebi.ac.uk/gxa/experiments/E-MTAB-970.
